# Transient telomerase mRNA delivery extends the proliferative capacity of patient-derived Duchenne muscular dystrophy myogenic progenitors

**DOI:** 10.64898/2026.11.25.769108

**Authors:** L. Barone, F Mourkioti, G Cossu

## Abstract

Duchenne muscular dystrophy (DMD) is an X-linked disease, characterised by progressive skeletal and cardiac muscle degeneration and impaired regeneration. Repeated regenerative cycles contribute to muscle stem cell exhaustion, potentially driven by telomere shortening under chronic proliferative demand. Here, we tested whether transient delivery of hTERT mRNA can increase the relative telomere signal (Q-FISH) in patient-derived DMD myogenic progenitors (CD56+) and interstitial (CD56−) cells from two DMD donors, compared with one control donor. hTERT mRNA treatment increased the relative telomere signal and added 2 to 6 population doublings in DMD myogenic progenitors, whereas healthy donor cells showed no comparable benefit in proliferation. Exploratory transcriptomic analysis indicated genotype-associated differences in translation- and cell-cycle-related gene sets following hTERT treatment. Telomere elongation and improved proliferation were not accompanied by recovery of myogenic differentiation; no karyotypic abnormalities were detected under the conditions tested. By transiently increasing the proliferative output of dystrophic cells without genomic integration, this proof of concept provides a basis for further investigation of telomere-associated proliferative limits in DMD and a starting point for future regenerative approaches.

## Introduction

Chronic muscle degeneration in Duchenne muscular dystrophy (DMD) places an immense proliferative burden on resident stem cells. Caused by loss-of-function mutations in the DMD gene, the absence of dystrophin compromises myofibre integrity and triggers persistent tissue damage (Duan et al, 2021). Following acute injury in healthy tissue, satellite cells, the resident muscle stem cells, activate, proliferate, and fuse to regenerate myofibres, while re-establishing a reservoir of quiescent satellite cells to preserve tissue homeostasis (Zammit et al, 2004). In DMD, however, unrelenting degeneration forces satellite cells into continuous rounds of proliferation. Over time, this chronic demand impairs their capacity to expand and regenerate, favouring fibroadipose substitution, progressive muscle weakness, and eventual cardiorespiratory failure (Chulanova et al, 2025).

This repeated activation imposes a sustained proliferative demand on satellite cells and their myogenic progeny. Each regenerative cycle requires expansion of the progenitor pool, increasing the number of cell divisions experienced by these cells (Blau et al, 1983; Mouly et al, 2005). In the inflammatory and metabolically demanding environment of DMD muscle, additional stressors may further accelerate telomere erosion, progressively limiting the replicative potential of muscle progenitors (Consalvi et al, 2011; Shalev et al, 2013). Although several studies have shown that non-canonical myogenic cells, such as interstitial, mesenchymal and mesodermal cells, contribute to muscle regeneration, none of them can fully regenerate muscle (Peault et al, 2007).

Because every round of cell division results in progressive telomere shortening, tissues exposed to chronic regenerative demand are uniquely vulnerable to critical telomere attrition (Mouly et al, 2005). As capping structures protecting chromosome ends, functional telomeres prevent DNA damage response (DDR) activation and cell-cycle arrest (Shay & Wright, 2011). Importantly, even a single critically short telomere can become dysfunctional (Hemann et al, 2001), triggering a persistent DDR cascade that blocks cell-cycle progression to prevent end-to-end chromosome fusions and genomic instability (d’Adda di Fagagna et al, 2003). To counter this attrition, cells rely on telomerase; however, telomerase expression is widely repressed in most adult tissues, with detectable activity restricted to a few highly proliferative compartments (Wright et al, 1996).

Seminal work by Helen Blau and colleagues first described the limited replicative lifespan of DMD myoblasts (Blau et al, 1983; Webster & Blau, 1990). Building on this foundation, subsequent studies demonstrated how critical telomere shortening drives cell fate decisions and exacerbates pathology across different tissues (Sacco et al, 2010; Mourkioti et al, 2013; Tichy et al, 2017, 2021). The risk that myogenic progenitors reach the Hayflick limit prematurely raises the question of whether restoring telomere length can prevent their proliferative exhaustion, and whether telomerase activity can sustain long-term stem cell capacity under the unrelenting stress of DMD.

Telomerase discovery enabled in vitro immortalisation by hTERT overexpression (Bodnar et al, 1998), mimicking the mechanism by which most cancers evade replicative senescence (Shay & Wright, 2011). Over the past 30 years, several cell types have been successfully transduced with telomerase, achieving virtually unlimited proliferation and providing a powerful research tool, but raising safety concerns for therapeutic applications (Shay & Wright, 2011).

The recent rise of mRNA-based technologies has highlighted their potential to provide transient and tunable protein expression without permanent genomic modification (Pardi et al, 2018). In this context, transient expression of telomerase could offer a safer alternative to constitutive or inducible hTERT expression (Zhu et al, 2007), allowing telomere restoration while preserving the physiological control of cellular proliferation. The delivery of hTERT-encoding mRNA, first reported in human fibroblasts and myoblasts (Ramunas et al, 2015), has since been applied to HGPS fibroblasts (Li et al, 2019) and human skin wound healing (Chang et al, 2024). In that first study, three transfections extended telomeres and conferred +28 ± 1.5 population doublings (PDs) in fibroblasts but only +3.4 ± 0.4 PDs in myoblasts, so the gain in the myogenic precursors most relevant for regenerative medicine was modest. Whether this approach can improve the proliferative capacity of dystrophic myogenic cells has not been tested.

Here, we tested transient hTERT mRNA delivery in donor-derived DMD myogenic progenitors (CD56+) and interstitial cells (CD56−) compared with a control donor. Despite enzyme production in all populations, an increase in proliferative output was observed only in DMD cells, and telomere elongation was most evident in DMD cells, suggesting a possible disease-associated response. DMD cells had shorter telomeres than control cells at baseline, and transient hTERT expression was associated with additional population doublings in DMD-derived cultures without detectable karyotypic abnormalities, but also without recovery of myogenic differentiation. Exploratory transcriptomic profiling further indicated a genotype-dependent response to treatment.

## Results

### DMD-derived myogenic progenitors display impaired proliferative and differentiation potential at baseline

We first established the baseline characteristics of the isolated cell populations. Cells derived from 3 different donors (2 DMD patients, aged 18 and 14 years, and 1 control, aged 9 years) were separated by magnetic beads and analysed by flow cytometry (Fig. 1A). The gating strategy is shown in Fig. S1. Healthy cells showed a higher proportion of CD56+ cells than both DMD donors (35.2% vs 16.9% in DMD #1 and 6.9% in DMD #2 in the unsorted populations; Fig. 1A). After magnetic separation, the CD56+ fractions contained 95.8% (control), 78.0% (DMD #1) and 10.0% (DMD #2) CD56+ cells, while the residual CD56+ cells in the CD56− fractions were 6.7%, 7.1% and 2.3%, respectively (Fig. 1A). While CD56+ cells are enriched in myogenic progenitors, mostly satellite cells (Meng et al, 2011), CD56− cells represent a heterogeneous population of interstitial cells, which we did not separate further to avoid additional stress to the cells. Throughout, we refer to the CD56+ and CD56− fractions as myogenic progenitors and interstitial cells; these terms reflect the selection strategy rather than the purity of each fraction. Notably, magnetic separation enriched CD56+ cells efficiently in DMD #1 but not in DMD #2, where the nominal CD56+ fraction contained only 10% CD56+ cells. Accordingly, results from the DMD #2 CD56+ fraction cannot be attributed specifically to myogenic progenitors. CD56− cells made up the majority of all unsorted populations, and an even larger share in the DMD samples (92.3% in DMD #2; Fig. 1A). The low CD56+ content of the DMD samples, most marked in DMD #2, is consistent with progressive depletion of the satellite cell pool in dystrophic muscle (Decary et al, 2000).

**Figure 1).**
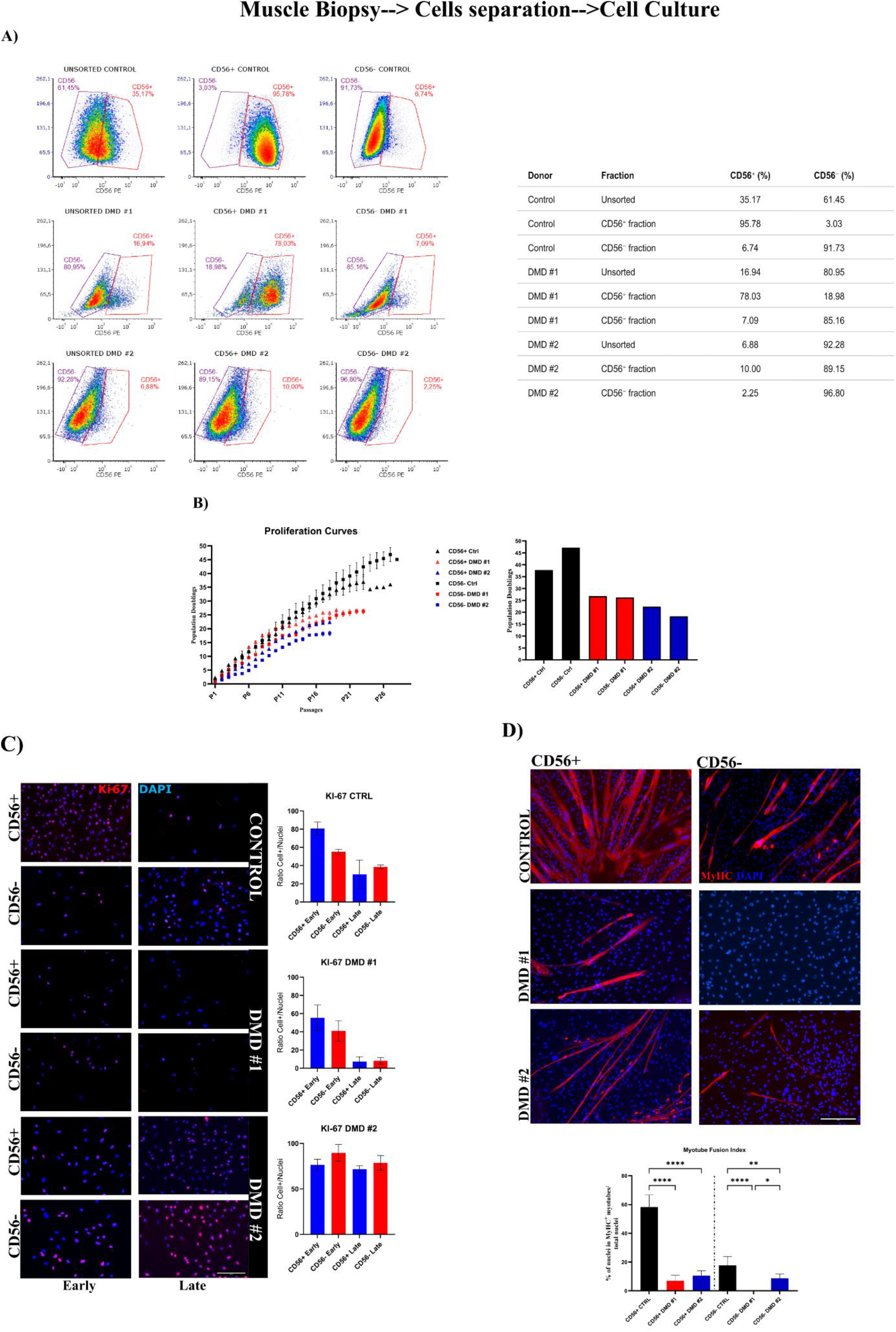
Flow cytometry analysis of myogenic cells. A) Analysis of healthy (left), DMD #1 (centre) and DMD #2 (right) cells extracted from biopsies: The first lane shows the original cell composition (unsorted) after a few passages in culture. The second and third lanes show the cell composition after magnetic bead separation. Data were acquired in three different experiments. The table on the right summarises the percentages for all donors. B) Proliferation curves of myogenic progenitor cells from a healthy donor (black) and two DMD donors (red and blue). The x axis shows the passages in culture. Cells were seeded at equal density (8 × 10⁴) at each passage to calculate population doublings (PD; harvested cells = seeded cells × 2^PD). Curves from two parallel cultures derived from the same sorted population (technical duplicates) are shown; passages (P) are counted from the magnetic separation (P1). On the right, summary plot of the final PD reached. Data are shown as mean ± SEM. C) Immunofluorescence staining for Ki-67 (red) and DAPI (blue) to evaluate cells in active division at different stages in each donor. Passages were matched as closely as possible between populations according to their stage of ageing. The percentage of Ki-67-positive cells was calculated over the total number of nuclei. Data represent mean ± SD of at least three technical replicates from every donor. At least 200 nuclei per condition were analysed. No statistical test was applied. Scale bar: 300 µm. D) Immunostaining for MyHC to assess differentiation of CD56+ and CD56− cells from dystrophic and healthy donors, stage-matched (early time point) to establish baseline differentiation potential. The panel on the right shows the myotube fusion index. Data were analysed by one-way ANOVA followed by Šídák’s post-hoc test. *p < 0.05, **p < 0.01, ****p < 0.0001. Scale bar: 500 µm.

After separation, cell populations were cultured in parallel in a newly devised, in-house proliferation medium until replicative exhaustion, to determine their proliferative potential. Proliferation was measured as population doublings (PD). Control CD56+ and CD56− cells reached 38 and 47 PD, respectively (mean of two cultures), compared with 27 (CD56+) and 26 (CD56−) in DMD #1 and 22 (CD56+) and 18 (CD56−) in DMD #2 (Fig. 1B and Table 1). No major morphological differences associated with cellular ageing were observed between control- and DMD-derived cells during culture (Fig. S2). At early and late passages we stained the cells for Ki-67 to see how many were cycling (Fig. 1C; values in Table S1). Control cells had more Ki-67 positive cells than DMD #1 cells at both time points. DMD #2 was unexpected: most of these cells were Ki-67 positive at both time points, although it was the donor that proliferated the least. This could mean that these cells move slowly through the cell cycle and keep Ki-67 expressed, which fits with the low EdU labelling shown later. We also stained for senescence-associated β-galactosidase (SA-β-gal), which was prominent in control cultures and weak or absent in DMD cultures (Fig. S3).

**Table 1.).**
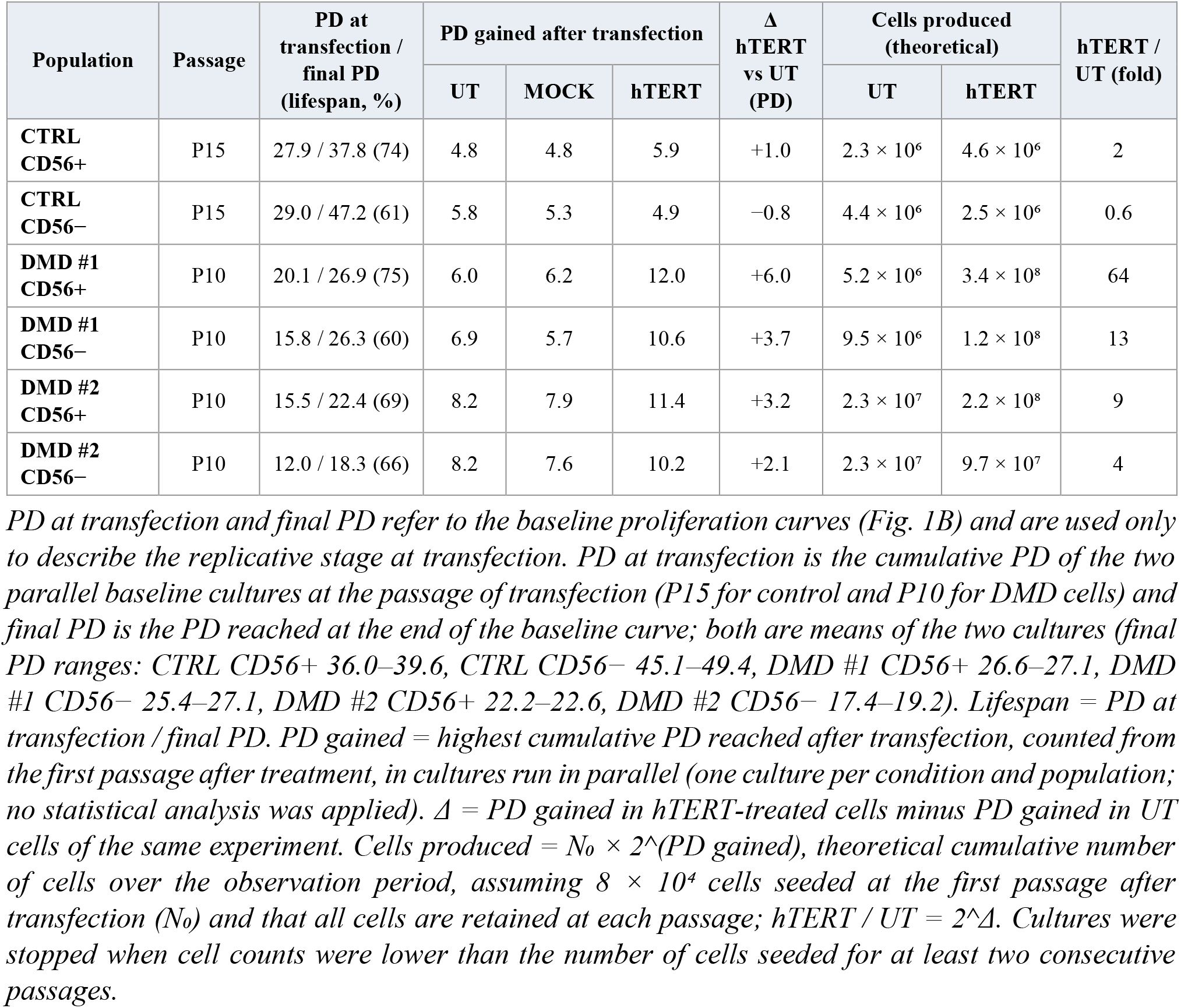
Population doublings (PD) at transfection and gained after hTERT mRNA treatment, and theoretical cell production, in untreated (UT), mock-transfected (MOCK) and hTERT-treated cultures.

To evaluate differentiation, cells were cultured in low-serum differentiation medium and assessed for MyHC expression and fusion index (percentage of nuclei within MyHC+ myotubes; Fig. 1D). Control cells served as an internal control of differentiation, and even the CD56− population showed some MyHC expression (CTRL CD56+ 58.3%, CD56− 17.7%). DMD cells differentiated to a limited extent (DMD #1 CD56+ 7.0%, CD56− 0%; DMD #2 CD56+ 10.5%, CD56− 8.7%), and even these low levels were lost in the following passages, highlighting reduced differentiation in DMD myogenic progenitors at late stages (Webster et al, 1986).

### DMD cells exhibit reduced telomere length compared to healthy controls

Given the reduced proliferative capacity observed in DMD progenitors, we next investigated whether differences in telomere length and telomerase activity were present at baseline. To establish a baseline, the relative telomere signal was measured at single-cell resolution by Q-FISH at the start of culture (Fig. 2A). The DMD-derived cultures showed lower relative telomere signal than controls, with the most pronounced shift in telomere distribution observed in the CD56+ compartment (Fig. 2B). In the CD56− population the profile was more homogeneous, and a within-experiment nuclear-level comparison indicated a difference from control for DMD #2, but not for DMD #1.

**Figure 2).**
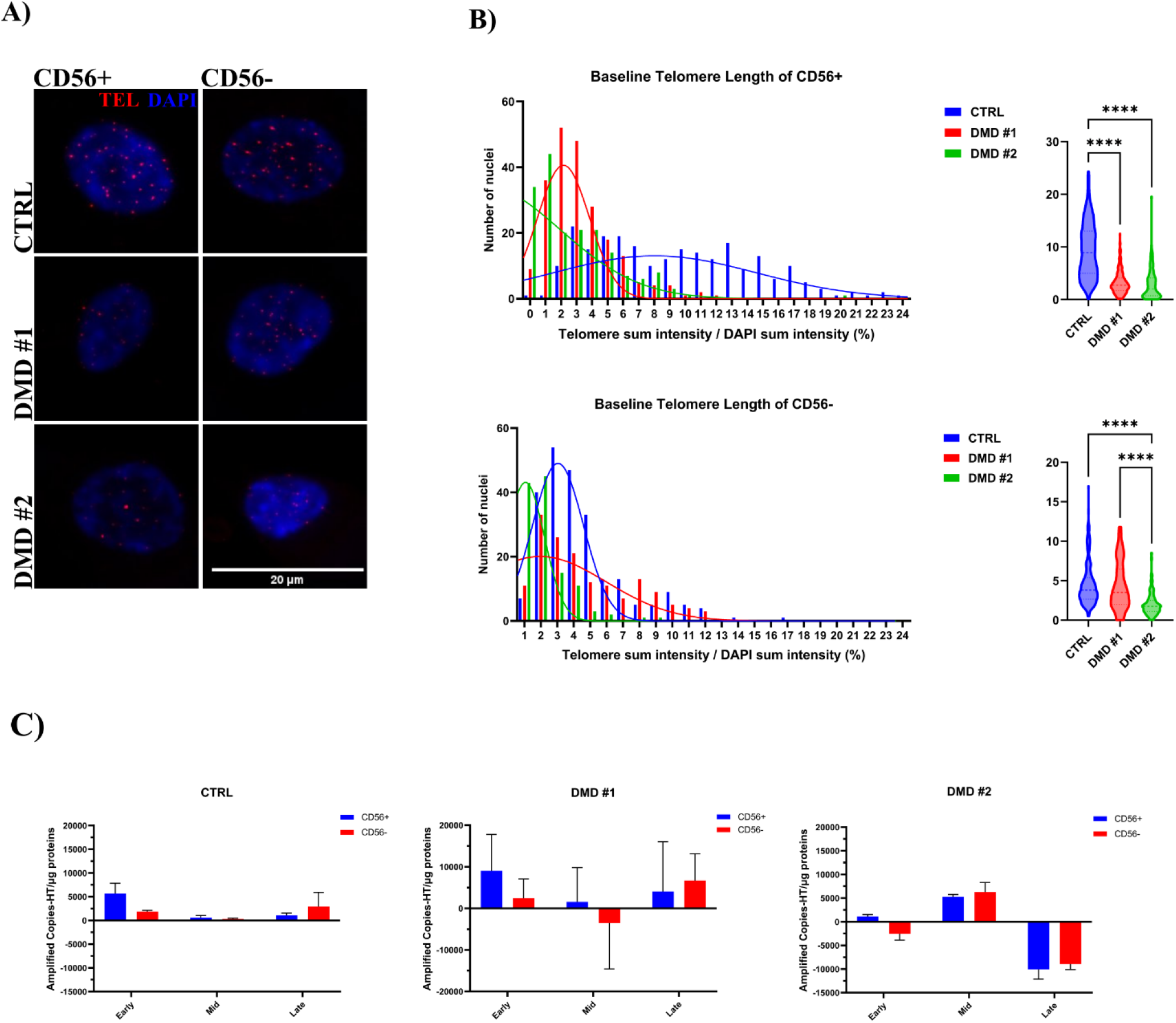
A) Representative images of Q-FISH-stained nuclei and their quantification (on the side). Myogenic cells from the three donors were stained to quantify telomere length (red) at the early stage of culture. B) Frequency distribution of telomere intensity in single cells (bar plot, left) and overall distribution (violin plot, right) for each population (control in blue, DMD #1 in red, DMD #2 in green). The y axis shows the number of nuclei evaluated and the x axis the normalised telomere intensity. Statistical significance was assessed by one-way ANOVA followed by Tukey’s multiple comparisons test, with nuclei as the unit of analysis within a single experiment per donor. Data are presented as mean ± SEM for ANOVA comparisons. Scale bar: 20 µm. ****p < 0.0001. C) Telomerase activity (TRAP) at early, mid and late culture stages. The y axis shows amplified copies minus heat-treated (HT) control per µg of protein. Bars: mean ± SD of three technical replicates of a single extract. Negative values indicate that the heat-inactivated control exceeded the sample signal, and are interpreted as activity below the detection limit. No statistical comparison was applied given the technical nature of the replicates and the variability of the assay.

To assess endogenous telomerase activity during in vitro expansion, TRAP assays were performed on extracts collected at early, mid and late stages of culture (Fig. 2C; stages defined in Methods). In control cells, activity was detectable at the early stage and, in the CD56+ population, decreased approximately tenfold by the mid stage. In DMD cultures, activity was variable between stages and donors, ranging from levels comparable to or higher than control (DMD #1 early; DMD #2 mid) to values below background (DMD #2 late). Positive controls (Fig. S4) confirmed assay performance. Thus, endogenous telomerase activity was detectable in almost all cultures but was not enough to sustain proliferation in DMD cells, which combined shorter telomeres (Fig. 2B) with a limited lifespan. This provided the rationale for supplying exogenous hTERT.

### hTERT mRNA transfection produces functional telomerase and increases proliferative output in DMD-derived cells

We next tested whether transient hTERT expression could improve the proliferative capacity of DMD progenitors. The size and integrity of the *in vitro*-transcribed mRNA encoding hTERT (Fig. 3A) were confirmed by capillary electrophoresis using TapeStation (Fig. S5). Transfection conditions were optimised using T7 GFP-reporter mRNA in HeLa and control cells (Fig. S6). Following hTERT transfection, hTERT immunoreactivity was higher in all transfected populations than in untreated cells (Fig. 3B). The increase was largest in DMD-derived progenitors, up to approximately 6.8-fold (DMD #2 CD56+), and was statistically significant in the four DMD populations but not in the control ones. To confirm the functional activity of the expressed protein, protein extracts from the same cells were analysed by TRAP 24 h after transfection (Fig. 3C). Telomerase activity was higher in all hTERT-treated samples, significantly so in every population except DMD #2 CD56−, which confirms delivery and translation of the mRNA and the enzymatic activity of the encoded protein. To assess the transient nature of hTERT expression, cells were immunostained again one week after transfection. At this time point, no differences among experimental groups were detected (Fig. S7), supporting the transient expression of the transfected mRNA.

**Figure 3).**
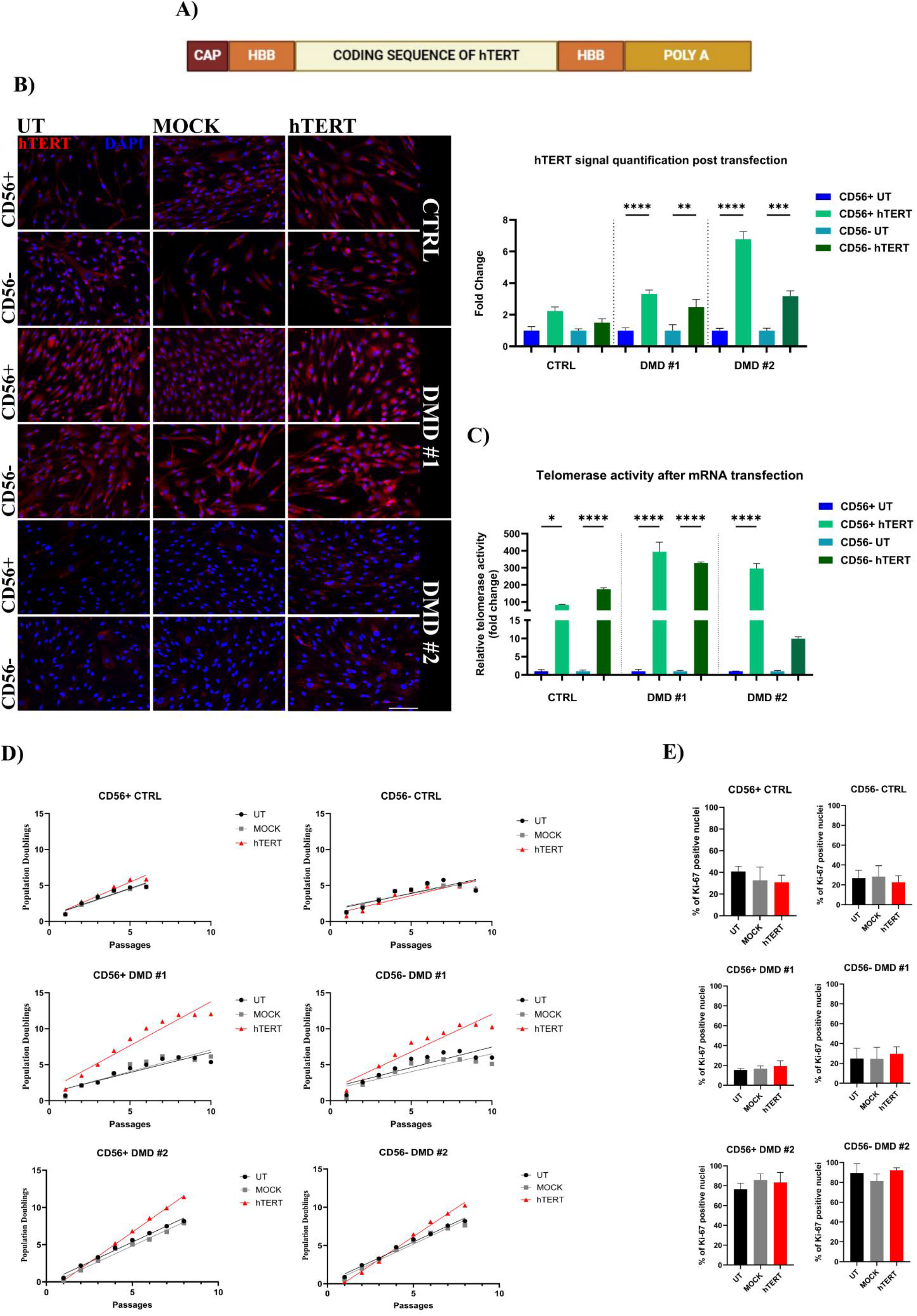
Efficiency assessment of hTERT mRNA transfection at 1 µg/ml. A) Schematic structure of hTERT mRNA sequence synthesised by IVT (in vitro transcription). B) hTERT expression after hTERT mRNA transfection (1 µg/ml). Immunofluorescence for hTERT (red) and nuclei (DAPI, blue) in CD56+ and CD56− muscle progenitors from healthy (CTRL) and DMD donors. Columns correspond to untreated (UT), mock-transfected (MOCK), and hTERT mRNA-transfected samples (hTERT). Rows represent CD56+ and CD56− populations for each donor. Cells were fixed 48 h post-transfection. Stronger nuclear and cytoplasmic hTERT staining is observed in transfected cells, especially in DMD-derived progenitors. Quantification of fluorescence intensity (bar plot on the right) is shown as fold change relative to the untransfected samples (CTRL CD56+ 2.23, CD56− 1.49; DMD #1 CD56+ 3.32, CD56− 2.48; DMD #2 CD56+ 6.78, CD56− 3.18) and analysed by one-way ANOVA with Šídák’s multiple comparisons post-hoc test on at least 3 independent fields per condition. Mock-transfected samples were quantified only for the control and DMD #1 donors, are not shown in the plot and did not differ significantly from untreated samples. Scale bar: 300 µm. **p < 0.01, ***p < 0.001, ****p < 0.0001. C) Telomerase activity quantification after hTERT transfection. Cell proteins from the three donors were harvested 24 h after hTERT transfection. Values are displayed as fold change relative to the untreated counterpart. Data were normalised as described in Methods and analysed by one-way ANOVA (comparing only treated and untreated data of each sample) with Šídák’s multiple comparisons post-hoc test. Error bars: mean ± SEM. *p < 0.05, ****p < 0.0001. D) Evaluation of proliferation after hTERT mRNA transfection (1 µg/ml) in muscle progenitors. Cells were treated at passage 15 (CTRL) or passage 10 (DMD-derived cells), according to their replicative stage; the x axis indicates passages after transfection, and the lines are simple linear regressions. One culture per condition and population was analysed; therefore, no statistical analysis was applied. Cultures were stopped at the last point shown, after at least two consecutive passages yielding fewer cells than those seeded. E) Proliferative marker evaluation after hTERT mRNA transfection. Representative images of Ki-67 (red) and DAPI (blue) staining in myogenic cells two passages after transfection. Quantification of Ki-67+ cells was performed by counting positive nuclei over the total number of nuclei (expressed as percentage). At least 5 fields per condition were acquired. Data were analysed by one-way ANOVA followed by Dunnett’s multiple comparison test; no comparison was significant. Scale bar: 300 µm.

We next asked whether transient hTERT expression translated into a functional change in proliferative capacity (Fig. 3D and Table 1). Transfection was performed after substantial replicative expansion of each population (passage 15 for CTRL and passage 10 for DMD-derived cells), corresponding to approximately 61–74% (CTRL) and 60–75% (DMD) of the final population doublings that each population reached in the baseline curves (Table 1). While control cells showed no differences between untreated, mock-transfected, and hTERT-treated conditions, both DMD donor-derived cultures showed additional population doublings following hTERT treatment. The effect was most pronounced in DMD #1, which showed the largest increase in proliferative output. These observations suggest a treatment-associated increase in proliferative output in the DMD-derived cultures tested, but the post-transfection curves reflect one culture per condition and cannot establish reproducibility or a disease-specific effect.

To check whether hTERT changed cell-cycle activity, we measured Ki-67 two passages after transfection (Fig. 3E). There were no significant differences among the groups. However, Ki-67 stayed high in DMD #2 despite its low proliferation rate, so in this donor we also measured DNA synthesis, co-staining Ki-67 and EdU after a 2-h pulse at the same time point (Fig. S8). Across populations and conditions, 76.4–92.2% of the nuclei were Ki-67 positive, but only 24.0–39.5% had incorporated EdU, and hTERT had no significant effect on EdU incorporation. In these cultures, therefore, Ki-67 positivity did not directly reflect DNA synthesis. Since EdU incorporation did not differ between conditions, it does not explain the gain in population doublings after treatment.

### hTERT mRNA transfection led to telomere elongation with no detectable karyotypic abnormalities

To determine whether the proliferative improvement induced by transient hTERT expression was accompanied by changes in telomere biology, the relative telomere signal was quantified at single-cell resolution by Q-FISH two passages after transfection (Fig. 4A). Compared with untreated cells, hTERT-treated cultures displayed higher telomere signals in within-experiment comparisons of nuclei from both DMD donors (CD56+ and CD56−), more markedly in DMD #2 than in DMD #1 (Fig. S10). In the CD56− control population, an increase of similar size was also observed in mock-transfected cells (Fig. S9–S10) and therefore cannot be attributed to hTERT. In contrast, CD56+ control cells did not show an increase in relative telomere signal and instead exhibited a reduction during the same time window.

**Figure 4).**
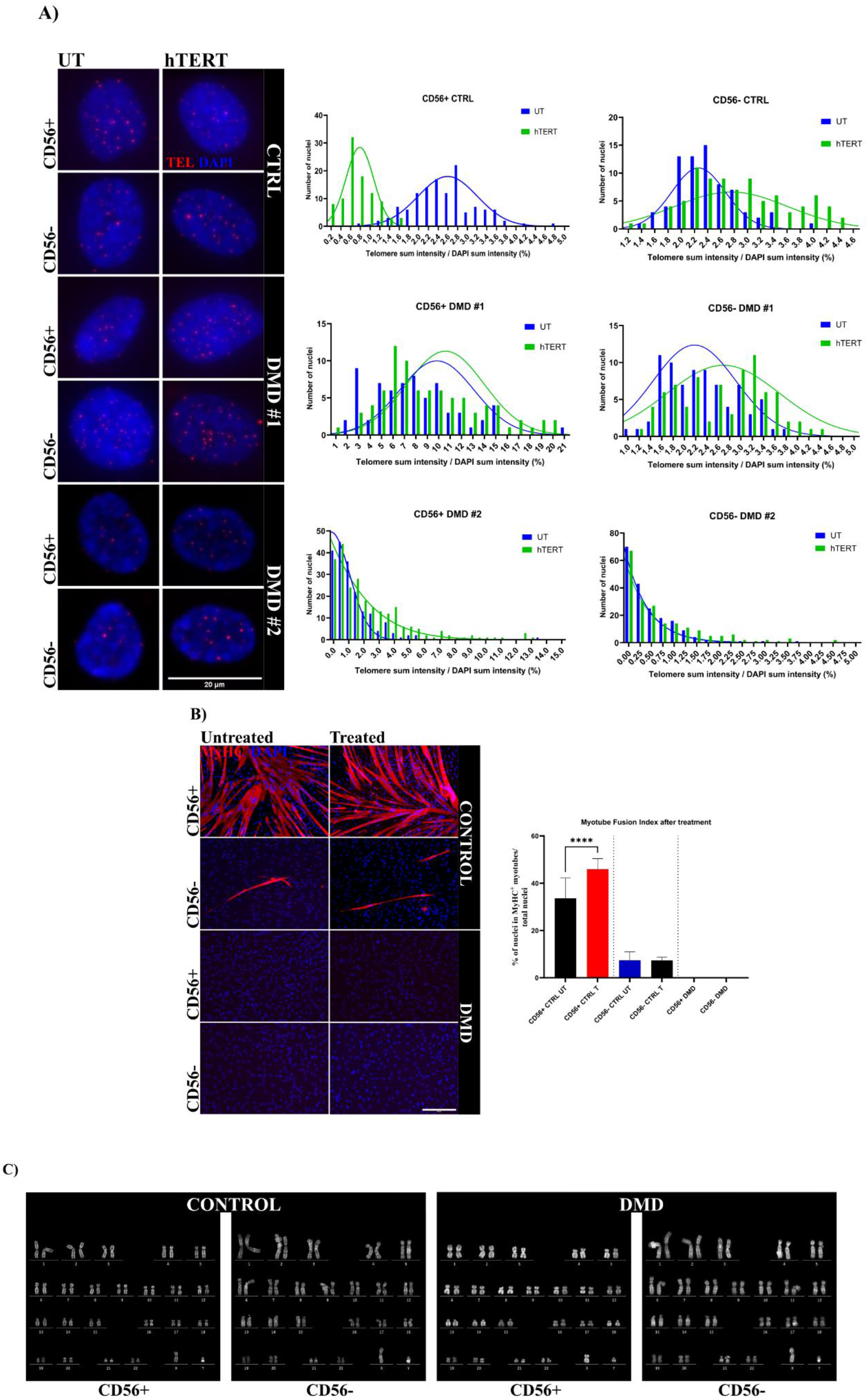
A) Representative images of Q-FISH-stained nuclei in myogenic cells (left) and relative telomere signal after treatment with hTERT-encoding mRNA (right): frequency distribution of telomere fluorescence intensities in CD56+ and CD56− cells from control (CTRL) and Duchenne muscular dystrophy (DMD) donors. Histograms represent individual nuclear measurements (bins defined automatically) of untreated (UT, blue) and treated (T, green) cells, overlaid with Gaussian-fit curves. Scale bar: 20 µm. B) Representative images of the differentiation assay after treatment, with the myotube fusion index on the side. Cells were induced to differentiate approximately one week after hTERT mRNA transfection. Images show immunostaining for MyHC (red) and nuclei (blue) in the CD56+ and CD56− populations of control and DMD cells (DMD #1 shown as representative; no MyHC-positive cells were detected in either DMD donor or population, before or after treatment). The bar plot shows the fusion index (see Fig. 1D). Results were analysed by one-way ANOVA followed by Šídák’s post-hoc test. Error bars: mean ± SEM. ****p < 0.0001. Scale bar: 500 µm. C) Karyotype images of late-passage control cells and late-passage hTERT-treated DMD cells (CD56+ and CD56−). Cells were analysed by the Cytogenetics Laboratory of San Raffaele Hospital.

Beyond changes in the mean telomere signal, single-cell analysis revealed a shift of the relative telomere signal distribution toward higher values following hTERT treatment in DMD-derived progenitors. This effect was particularly evident in the CD56+ population, where the proportion of nuclei with lower telomere signals decreased after treatment (bar plot in Fig. 4A). The CD56− population displayed a more homogeneous distribution, although a similar trend toward telomere elongation was observed.

Although DMD #2 CD56− cells did not reach statistical significance in the TRAP assay (Fig. 3C), Q-FISH nevertheless showed a modest increase in relative telomere signal in this population. We then tested the differentiation of treated cells: in controls, the fusion index of CD56+ cells was significantly higher than in untreated cells, whereas CD56− cells showed no difference. In both DMD donor-derived CD56+ and CD56− populations, no MyHC-positive cells were detected at this passage, either before or after treatment (DMD #1 shown as representative; Fig. 4B), so this assay did not demonstrate preservation or restoration of myogenic differentiation in the DMD-derived cultures. Finally, we assessed chromosomal stability by karyotype analysis of late-passage control cells and late-passage hTERT-treated DMD cells (at least 20 metaphases per sample), which showed a normal karyotype (Fig. 4C).

Together, these results show that transient hTERT mRNA delivery increases the relative telomere signal in the populations studied, most clearly in DMD-derived cultures, without detectable chromosomal abnormalities under the conditions tested.

### Transcriptional signatures associated with genotype and hTERT treatment

To explore the molecular programmes associated with the DMD phenotype and with the response to hTERT mRNA, we performed bulk RNA-sequencing one week after treatment, followed by gene set enrichment analysis (GSEA). Because the dataset contains one control donor and two DMD donors, with one sample per donor and condition and no independent biological replication, all RNA-seq contrasts are exploratory and descriptive. We compared DMD with control cells in the untreated condition (baseline) and examined the genotype × treatment interaction, which was used to explore possible differences in the hTERT-associated transcriptional response between the sampled DMD and control cells (Fig. 5).

**Figure 5).**
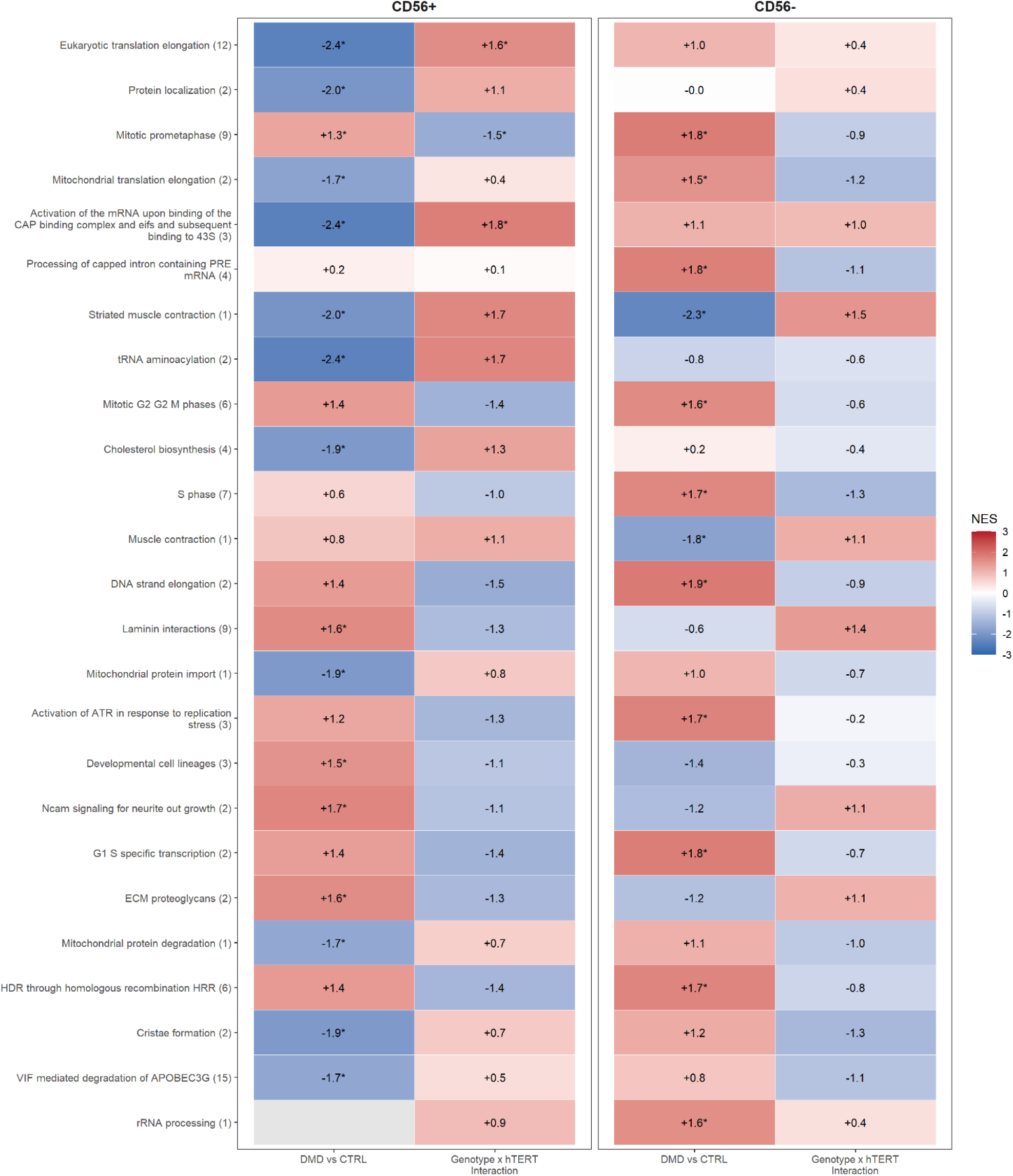
Reactome pathway clusters in CD56+ and CD56− cells. Heatmaps show the mean normalised enrichment score (NES) of Reactome pathway clusters (number of pathways in parentheses) for DMD vs control in untreated cells (DMD vs CTRL) and for the genotype × hTERT interaction. Pathways with padj < 0.25 in at least one contrast were grouped by the overlap of their leading-edge genes (Jaccard ≥ 0.3), and the 25 clusters with the most significant pathways are shown; each cluster is named after its most significant pathway. Red, positive NES; blue, negative NES. For the interaction, a positive NES indicates that the effect of hTERT is more positive in DMD than in control cells. Asterisks indicate at least one pathway of the cluster with padj < 0.05; grey, pathways not tested. One control and two DMD donors, no replicates.

Among CD56+ cells, DMD cells showed lower enrichment of clusters related to translation (elongation, cap-dependent initiation and tRNA aminoacylation), mitochondrial function (translation, protein import and degradation, cristae formation), cholesterol biosynthesis and striated muscle contraction, and higher enrichment of clusters related to mitosis (prometaphase), laminin interactions, extracellular matrix proteoglycans and developmental lineages (Fig. 5). Clusters related to G2/M, DNA replication and DNA repair showed positive enrichment that did not reach the significance threshold. Among CD56− cells, DMD cells showed higher enrichment of clusters related to cell-cycle progression and DNA replication and repair, including ATR activation in response to replication stress and homologous recombination, and lower enrichment of muscle contraction clusters (Fig. 5). This is counterintuitive, since DMD cells proliferate less, and we come back to it in the Discussion. Translation and mitochondrial clusters changed in opposite directions in the two populations, being lower in CD56+ and higher in CD56− cells. Hallmark gene sets showed a consistent picture for cell-cycle programmes (E2F targets, G2M checkpoint and mitotic spindle, higher in DMD cells in both populations; Fig. S11).

In the interaction, only the translation elongation (positive) and mitotic prometaphase (negative) clusters in CD56+ cells crossed the nominal analysis threshold, and no cluster did so in CD56− cells. For most clusters, however, the interaction had the opposite sign to the baseline difference in both populations: clusters related to mitosis, DNA replication and repair, which were higher in DMD cells at baseline, were lower in the interaction, and translation and striated muscle contraction clusters, which were lower at baseline in CD56+ cells, were higher. In Hallmark gene sets, E2F targets, G2M checkpoint and mitotic spindle were also significantly lower in the interaction in CD56+ cells (Fig. S11), whereas the interferon alpha response did not reach significance in either contrast. Because the baseline and interaction contrasts share the untreated control sample, part of this opposition is expected, and as most of these changes were not significant we consider it a trend requiring validation and not a finding (pathway-level results in Fig. S12–S13).

## Discussion

### Transient hTERT extends the replicative lifespan of DMD progenitors

It has been debated whether DMD progression depends on replicative exhaustion of satellite cells or impaired differentiation (Oexle & Kohlschütter, 2001). Our results are consistent with a telomere-associated proliferative limitation in the studied DMD-derived cultures that may be partially relieved by transient hTERT expression: a single transfection of hTERT mRNA was associated with approximately 2 to 6 additional population doublings in DMD-derived cultures, corresponding to a theoretical 4- to 64-fold increase in cell production under the assumptions of Table 1, while control cells showed no consistent gain (Table 1). This observation may be relevant to cell-expansion strategies, although therapeutic myogenic function was not preserved in these experiments. In the first trial of HLA-matched donor mesoangioblasts in DMD, which was feasible and relatively safe, engraftment was minimal and the authors argued that the cell dose had not approached the level needed for efficacy (Cossu et al, 2015). Autologous cells avoid rejection and can be genetically corrected, but they must be expanded from a small biopsy, and DMD progenitors stop dividing early (Webster & Blau, 1990).

These results move beyond the work of Ramunas et al (2015), who showed that modified TERT mRNA lengthens telomeres and adds doublings in healthy human fibroblasts and myoblasts without immortalising them. They noted that DMD myoblasts have short telomeres and a limited replicative capacity (Webster & Blau, 1990; Sacco et al, 2010; Mourkioti et al, 2013) and proposed the approach for disease modelling, drug screening and regenerative medicine, but used only healthy cells. In human DMD cells the link between short telomeres and a limited lifespan has been correlative, and causal evidence comes from the mouse, where short telomeres worsen the dystrophic phenotype (Sacco et al, 2010).

Transient hTERT expression in DMD myogenic progenitors was associated with additional population doublings after one transfection of unmodified mRNA, whereas the earlier study used three doses of modified, less immunogenic mRNA (Karikó et al, 2008). In DMD #1 CD56+ cells the gain (6.0 doublings) exceeded that of healthy myoblasts in that study (3.4), whereas in DMD #2 it was comparable (3.2), although the comparison is only indicative because lifespan depends on culture conditions: the myoblasts used in this study were grown under standard conditions and stopped expanding after about eight doublings, whereas ours were grown at 5% O2 in a proliferation promoting medium and the CD56+ controls reached 38 PD.

### Telomere extension accompanies the proliferative gain

Additional population doublings were accompanied by higher relative telomere signals, an association that does not by itself establish causality. In both DMD donors the telomere signal increased after treatment and fewer cells had lower telomere signals, mainly among CD56+ cells (Fig. 4A, S10), whereas control CD56+ cells, which showed no consistent gain, did not show increased relative telomere signals. Telomerase acts mostly on the shortest telomeres (Britt-Compton et al, 2009), and Li et al (2019) found that hTERT mRNA increased the proliferation of progeria fibroblasts with short telomeres but not of a line with normal telomere length. Our data follow the same pattern: at treatment the control cells had about 10 (CD56+) and 18 (CD56−) doublings ahead, against 6 to 10 for DMD cells (Table 1), and longer telomeres at baseline (Fig. 2B), so there was less for telomerase to repair. Delivery is unlikely to explain the difference on its own, since hTERT immunoreactivity rose more in DMD cells but telomerase activity increased in controls as well (Fig. 3B and 3C). Within each DMD donor, CD56+ cells gained more doublings than CD56− cells, in line with their larger telomere deficit at baseline (Fig. 2B), although the comparison is clearer in DMD #1, as the CD56+ fraction of DMD #2 was poorly enriched. Between donors the picture is less consistent: the increase in telomere signal was larger in DMD #2 than in DMD #1 (Fig. S10), yet DMD #2 gained fewer doublings (Table 1). DMD #2 had the shortest replicative lifespan of the three, reaching 22 (CD56+) and 18 (CD56−) population doublings against 27 and 26 in DMD #1. A larger relative gain in telomere signal therefore did not translate into proportionally more divisions, which is what would be expected if a telomere-independent barrier also limits these cells. Mock-transfected cultures behaved like untreated ones, gaining a comparable number of doublings in every population (Table 1) and, where telomeres were measured, showing no increase in telomere signal except in control CD56− cells (Fig. S9–S10). The proliferative gain therefore follows the mRNA rather than the transfection procedure. It does not follow that telomere elongation mediates it: TERT mRNA can reduce DNA damage and apoptosis in primary human cells without measurable telomere lengthening (Li et al, 2026), so without a catalytically inactive hTERT, elongation and additional doublings remain parallel consequences of treatment.

The proliferative association persisted beyond detectable exogenous hTERT expression. hTERT signal was back to baseline one week after transfection (Fig. S7), yet relative telomere signals remained higher at that time (two passages; Fig. 4A, S10) and the extra doublings accumulated over the following passages, consistent with a persistent effect after a brief pulse, although telomere elongation has not been established as the sole causal mechanism. Treated and untreated cultures of a donor stopped at about the same time, but the treated cells divided more in that time, so cumulative population doublings increased without a corresponding extension of time in culture. Thus, greater proliferative output should be distinguished from longer chronological culture lifespan. This suggests a second barrier that telomerase does not remove. Growth arrest through p16, which limits human myoblasts after TERT expression (Zhu et al, 2007) and increases as they become senescent (Francis et al, 2022), is a candidate, but we did not measure it. Late-passage DMD cultures were weak or negative for SA-β-gal (Fig. S3), but the stain is not specific for senescence (Gorgoulis et al, 2019) and lysosomal abnormalities described in dystrophic muscle (Jaber et al, 2025) could interfere with it, so we describe these cultures as replicatively exhausted and not as senescent.

### Telomere-associated proliferative limitations in DMD cells

In healthy adults most satellite cells are quiescent and divide only to repair damage (Relaix & Zammit, 2012), so their telomere reserve is used slowly, whereas in DMD continuous regeneration forces them to keep dividing. This agrees with the short telomeres and limited replicative capacity reported in dystrophic muscle cells (Decary et al, 2000; Sacco et al, 2010; Tichy et al, 2017) and with our baseline data: DMD progenitors had shorter telomeres than the control (Fig. 2B) and reached fewer doublings (Fig. 1B).

DMD cells did not divide faster than control cells in culture (Fig. 1B), so their shorter telomeres are unlikely to reflect a faster intrinsic division rate. In the patient, chronic regeneration calls satellite cells back to divide repeatedly, which likely causes replication stress. Exploratory RNA-seq patterns are compatible with this idea: DMD CD56− cells had higher DNA replication and replication stress sets (ATR activation, homologous recombination) than controls despite not dividing faster (Fig. 5), but with one control and no replicates we give it little weight. In DMD #2 most cells were Ki-67 positive but fewer incorporated EdU (Fig. S8), which we did not investigate further.

Age and disease stage have to be considered when reading these results. The control donor was 9 years old and the DMD donors 18 and 14, the biopsies came from different muscles, and all cells were measured after expansion, so we cannot tell how much of the difference between control and DMD cells comes from the disease, the age, the muscle or the donor. The same applies to the two DMD donors: DMD #2 had the shorter lifespan and the smaller gain although it was the younger of the two, so age does not account for the difference between them, and its nominal CD56 fraction contained only 10% CD56+ cells after sorting. Consequently, the response of this fraction cannot be assigned specifically to myogenic progenitors. Myoblast cultures can also be overrun by fibroblasts with passaging (Francis et al, 2022), so the composition of the CD56+ cultures may have changed during expansion. Age, disease stage and cell composition could all contribute, and two donors cannot separate them.

### Increased proliferative output does not restore myogenic differentiation

We also asked whether the extra proliferation would at least preserve the limited capacity of DMD cells to differentiate. DMD myogenic progenitors are known to differentiate less than healthy ones (Meng et al, 2018; Massenet et al, 2024), and in healthy human myoblasts fusion is also impaired at replicative senescence (Francis et al, 2022). Our DMD cells showed limited differentiation already at early passages and no differentiation a few passages later, with or without hTERT (Fig. 1D, 4B), so increased proliferative output did not preserve or restore differentiation under the conditions employed. The fusion index did not track CD56 enrichment: DMD #2, whose CD56+ fraction contained 10% CD56+ cells, reached a fusion index comparable to that of DMD #1, whose fraction was 78% pure (Fig. 1D). CD56 marks myogenic commitment but not the capacity to fuse, and in DMD the residual progenitor pool is itself functionally impaired (Meng et al, 2018; Massenet et al, 2024), so differentiation in these cultures is limited by the state of the cells rather than by the proportion of CD56+ cells in them. In control CD56+ cells, by contrast, the fusion index was higher after hTERT treatment (Fig. 4B). This was the only population in which treatment affected differentiation, and it was not seen in control CD56− cells. TERT mRNA has effects in primary human cells that do not require telomere lengthening (Li et al, 2026), which could act on differentiation independently of telomere length, but the observation comes from a single experiment and we did not test this. This distinction between expansion of muscle-derived cells and expansion of functionally myogenic progenitors is central to any therapeutic interpretation. The loss of myogenic differentiation has been reported in such cultures and we did not aim to restore it by other means such as expression of inducible MyoD.

### Limits and outlook

This is a proof of concept with the constraints of primary patient material: two DMD donors, a single control and cell banks expanded before they reached us, so earlier doublings are unknown. The post-transfection curves were based on one culture per condition, without independent biological replication or inferential testing; telomeres were evaluated at a single post-treatment time point using relative Q-FISH fluorescence, and we did not include a catalytically inactive hTERT control (Ramunas et al, 2015; Li et al, 2019). The RNA-seq has no independent biological replicates, cannot reliably separate donor and genotype effects, and may reflect secondary effects, because hTERT protein was back to baseline at one week (Fig. S7).

The effect might be increased with modified nucleotides, repeated doses, earlier treatment or more stable RNA formats, such as circular RNA (Wesselhoeft et al, 2018), and sorting the cells again before treatment would give purer populations. The mRNA approach avoids intentional genomic integration; treated cultures eventually stopped expanding and no abnormalities were detected in the karyotypes examined. These observations do not establish long-term genomic safety or exclude effects below the sensitivity of conventional karyotyping. A key test for any application is whether the expanded cells still regenerate muscle *in vivo*, which fusion in vitro cannot show.

### Conclusions

In patient-derived DMD progenitors, a single transfection of hTERT mRNA was associated with additional population doublings and higher relative telomere signals, whereas healthy cells, which were away from their proliferative limit and had longer telomeres, did not respond. These findings support further study of transient telomerase delivery as a means to investigate proliferative limits in DMD-derived cultures, but do not establish a causal telomere mechanism or disease-specific response. The gain was finite, as expected from a transient treatment, and did not extend to differentiation, which these cells already lacked. Our data offer an initial proof of concept for increasing the proliferative output of cells from DMD biopsies, without demonstrating that the expanded cells retain myogenic function, and point to what remains to be solved: a telomere-independent barrier to proliferation and the loss of myogenic function.

## Materials and methods

### Human samples and ethics

Skeletal muscle biopsies were obtained from one healthy donor and two patients with Duchenne muscular dystrophy (DMD #1 and DMD #2). The control was a 9-year-old male donor undergoing orthopaedic surgery at the Ospedale San Raffaele Scientific Institute; the biopsy was taken from the vastus medialis. Tissue procurement and all experimental procedures were approved by the Institutional Bioethical Committee of the Ospedale San Raffaele Scientific Institute (Protocol N°: UniMab), and written informed consent was obtained from the legal guardians before tissue collection, in accordance with the Declaration of Helsinki.

DMD #1 and DMD #2 were 18 and 14 years old, non-ambulant and not ventilated, and eligible for exon 51 skipping. They were enrolled in two Phase 1/2 trials of autologous human mesoangioblasts, DMD06-Mab (EudraCT number 2023-000148-4) and DMD04-Mab (EudraCT number 2019-001825-28), and their cells were obtained from an excision biopsy of the foot muscle extensor digitorum brevis (EDB). The protocols were approved by EMA and by the ethical committees of San Raffaele Hospital and the University of Manchester, respectively. The patients and their parents received an information sheet and signed an informed consent that included the donation of residual tissue to research.

### Cell culture and proliferation curves

Human muscle-derived cells were obtained from the biopsies of the three donors as described in Tonlorenzi et al (2007), and expanded to generate cell banks. DMD biopsies were processed in a GLP facility and expanded under GMP conditions, so the population doublings (PD) before our cultures are not available. All cultures were at passage 8 post-biopsy when plated in our laboratory, and passage numbers (P) in this study are counted from this point (P1). Cells were kept at 5% O2, 5% CO2 and 95% humidity in a proprietary medium based on low-glucose DMEM without phenol red, with 5% FBS, penicillin-streptomycin and a supplement mixture (composition protected by patent; the medium is available from the authors upon reasonable request). Cells were tested for mycoplasma every five passages and were always negative.

Cells were separated into CD56+ myogenic progenitors and CD56− interstitial cells (see below), as verified by flow cytometry, and each population was followed until natural exhaustion in vitro. For proliferation curves, 8 × 10⁴ cells were plated at every passage, harvested and counted manually in a Bürker chamber (cat. HS-1490). PD were calculated as PD = ln(harvested/seeded)/ln 2 and plotted cumulatively. Two parallel cultures derived from the same sorted population were followed for each population. Cultures were stopped when cell counts were lower than the number of cells seeded for at least two consecutive passages.

### Human TERT mRNA template generation and synthesis

A T7 plasmid containing a multiple cloning site, β-globin (HBB) UTRs and a poly-A sequence (pMRNAxp, mRNAExpress kit, cat. MR-KIT-1, Applied Bioscience) was used. The open reading frame of human TERT (NCBI NM_198253.2) was cloned into the vector by TwinHelix (The Netherlands), because the adjacent restriction sites prevented efficient digestion. The plasmid was amplified and the template was generated by PCR using the primer set supplied with the kit, following the manufacturer’s protocol that adds extra adenines, following the manufacturer’s protocol.

The fragment was run on a 0.8% agarose gel to confirm its molecular weight (approximately 3,340 bp), purified with Macherey-Nagel DNA spin columns (cat. 740609.50) and transcribed with HiScribe T7 (cat. E2065S) in the presence of an Anti-Reverse Cap Analogue (ARCA, Cap0), following the manufacturer’s instructions and using standard (unmodified) nucleotides. Residual DNA was removed with DNase. The transcript was purified by lithium chloride precipitation, air-dried, resuspended in RNase-free water and analysed with a TapeStation 4200 (Agilent, cat. G2991BA; Fig. S5). No further purification (for example HPLC or cellulose-based removal of double-stranded RNA) was performed.

### Magnetic bead separation

Cells expanded from the biopsies were harvested, resuspended in PBS with 1% FBS and 2 mM EDTA and filtered through a 70 µm strainer. After centrifugation and washing, cells were blocked and stained with anti-CD56 antibody-conjugated beads (Miltenyi, cat. 130-050-401) and loaded on an LD column (Miltenyi, cat. 130-042-901) mounted on a QuadroMACS separator (Miltenyi, cat. 130-091-051) in depletion mode. The flow-through contained the CD56− population; after removal of the column from the magnet, the retained CD56+ population was collected.

### Flow cytometry

Cells were harvested before and after magnetic separation, counted (at least 5 × 10⁴ cells), stained with a CD56-PE antibody (BD Pharmingen, cat. 555516) and analysed to determine the composition of the cell mixture. At least 3 × 10⁴ events were acquired per sample on a FACSCanto II (BD Biosciences) and analysed with FCS Express (De Novo Software).

### Cell treatment

Cells were plated in growth medium at 60–80% confluence and allowed to adhere overnight. The next day, the medium was replaced one hour before transfection, and hTERT mRNA was transfected with jetMESSENGER (Polyplus, cat. 101000005) following the manufacturer’s instructions. Briefly, the mRNA was mixed with jetMESSENGER buffer, spun down, combined with the transfection reagent, incubated for 15 min at room temperature, spun down again and added dropwise to the cells. The mRNA concentration was 1 µg/ml, and a single transfection was performed. Incubation times of 4 and 16 h (overnight) were tested, and 4 h was used for all further experiments. Mock-transfected cells received the transfection reagent without mRNA. Untreated (UT), mock-transfected and hTERT-treated cultures were run in parallel. Transfection conditions were optimised with a GFP reporter mRNA in HeLa and control cells (Fig. S6), and delivery of hTERT mRNA in DMD cells was verified by hTERT immunofluorescence and TRAP (Fig. 3B and 3C). Cells were transfected at passage 15 (CTRL) or passage 10 (DMD-derived cells), chosen according to the replicative stage of each population.

### Telomerase activity measurement

Telomerase activity was measured with the TRAPeze RT Telomerase Detection Kit (Merck, S7710) following the manufacturer’s protocol. Cell pellets were collected at the indicated time points, snap-frozen in liquid nitrogen and lysed in CHAPS buffer on ice. Protein concentration was determined by BCA assay (Thermo Fisher). Reactions contained 2 µL of protein extract (10 ng–1.5 µg of protein), the 5× TRAPeze RT reaction mix and Titanium Taq polymerase (Takara) in a final volume of 25 µL. Each run included a positive control, heat-inactivated extracts, a minus-telomerase control and a no-template control. Real-time amplification was performed on a StepOne system (Applied Biosystems) with the cycling conditions recommended by the manufacturer, and activity was expressed as amplified copies per µg of protein after subtraction of the heat-inactivated control. At baseline, early, mid and late stages of culture corresponded to approximately passage 2, 12 and 22 for control cells; for DMD #2, which has a shorter lifespan, the late stage corresponded to the last passages before proliferative arrest. After transfection, extracts were collected 24 h later and activity was expressed as fold change over the untreated cells of the same population.

### Immunofluorescence

Cells were washed with PBS, fixed in 4% paraformaldehyde (Sigma-Aldrich, 158127) for 15 min, rinsed with PBS, and blocked and permeabilised with 1% BSA and 0.2% Triton X-100 in PBS for 60 min. Cells were then incubated with primary antibodies for 1 h at room temperature or overnight at 4°C: anti-hTERT (rabbit polyclonal, Rockland, 600-401-252; the only anti-hTERT antibody tested that gave a specific signal, increased after hTERT mRNA transfection), anti-MyHC (MF20, Hybridoma Biobank) and anti-Ki-67 (SolA15, eBioscience, cat. 740008T). After PBS washes, cells were incubated with Alexa Fluor 488 or 594 secondary antibodies (Thermo Fisher) for 1 h at room temperature together with DAPI. Images were acquired with a Leica Eclipse Ts2 inverted fluorescence microscope and processed with ImageJ (Fiji).

### Ki-67 quantification

Ki-67 was quantified as the percentage of Ki-67-positive nuclei over the total number of DAPI-stained nuclei (at least 200 nuclei per condition).

### hTERT quantification

hTERT signal was quantified as the integrated density (IntDen) of each field normalised to the number of nuclei, using ImageJ, in at least three fields per condition (at least five in Fig. S7), and expressed as fold change over the untreated cells of the same population.

### EdU staining

DNA synthesis was assessed in DMD #2 cells one week (two passages) after transfection (4 h incubation). Cells were pulsed with 10 µM EdU for 2 h, fixed in 4% paraformaldehyde for 15 min, permeabilised, and EdU was detected with Click-iT Plus EdU Cell Proliferation Kit Alexa Fluor 594 (cat. C10639) following the manufacturer’s instructions. Cells were then stained for Ki-67 and DAPI as described above and imaged as in the immunofluorescence section. For each field, the percentage of Ki-67+EdU+ and Ki-67+EdU-nuclei was calculated over the total number of DAPI-positive nuclei, in 6–7 fields per condition from a single experiment.

### RNA extraction and sequencing

RNA was extracted with the RNeasy Mini Kit (Qiagen, cat. 74104) using columns with prior genomic DNA elimination, and its quality was checked with a TapeStation. Library preparation and sequencing were carried out by Novogene (Planegg, Germany). Messenger RNA was enriched from total RNA by poly-A selection with poly-T oligo-attached magnetic beads, fragmented, and reverse transcribed with random hexamer primers. Libraries were sequenced on an Illumina NovaSeq X Plus, generating 150 bp paired-end reads.

Preprocessing was performed by the provider. Raw reads were processed with fastp to remove adapter sequences, reads in which undetermined bases exceeded 10% and reads in which more than 50% of bases had a quality score of 5 or lower. Clean reads were aligned to the human reference genome (hg38) with HISAT2 v2.2.1 and gene-level counts were obtained with featureCounts v2.1.1, both with default parameters, so reads were counted without regard to strand. Libraries yielded between 31.6 and 38.1 million clean read pairs per sample, with Q30 above 96.8%, a total mapping rate of 98.2% to 98.5% and a unique mapping rate of 86.8% to 94.1%.

Downstream analysis of the gene-level count matrix was performed in R v4.6 using the DESeq2 package v1.52.0 (Love et al, 2014). The dataset comprised one control donor and two DMD donors, CD56+ and CD56− cells, untreated and hTERT-treated, with one sample per condition (12 samples in total), all sequenced in a single run. CD56+ and CD56− populations were analysed separately, after removing genes with fewer than 10 total counts. The design formula was ∼ Genotype + Treatment + Genotype:Treatment, with CTRL and untreated defined as the reference levels; the two DMD donors were treated as replicates of the DMD genotype and donor was not included in the model. The genotype effect (DMD versus CTRL) therefore refers to untreated cells, and the genotype by treatment interaction was used to determine whether the transcriptional response to treatment differed between DMD and CTRL samples. Statistical significance was assessed using Benjamini-Hochberg-adjusted p-values. Gene Set Enrichment Analysis (GSEA) was performed using the GSEA function implemented in the clusterProfiler package v4.20.2 (Yu et al, 2012). Genes were ranked according to their DESeq2 log2(fold changes), and enrichment analysis was conducted against Hallmark and Reactome gene set collections obtained from MSigDB (Liberzon et al, 2015) using the msigdbr package, with 10–500 genes per set. Ensembl gene identifiers were converted to gene symbols using the org.Hs.eg.db annotation package. Pathway enrichment was evaluated based on normalised enrichment scores (NES) and Benjamini–Hochberg-adjusted p-values. For visualisation, Reactome pathways with padj < 0.25 in at least one contrast were grouped by the overlap of their leading-edge genes (Jaccard index, average-linkage hierarchical clustering, clusters defined at a Jaccard index ≥ 0.3), and the 25 clusters containing the most significant pathways are shown; the score of each cluster is the mean NES of all its pathways, and an asterisk indicates at least one pathway with padj < 0.05. Given the number of donors and the absence of replicates, the analysis is exploratory and descriptive.

### Senescence-associated β-galactosidase assay

Senescence-associated β-galactosidase was detected with the Senescence β-Galactosidase Staining Kit (Cell Signaling Technology, cat. #9860) following the manufacturer’s instructions. Cells were washed, fixed in 4% paraformaldehyde for 15 min and washed in PBS, then incubated overnight at 37°C with the staining mixture (Staining Solution 10×, Reagent B, Reagent C, distilled water and X-gal solution pre-warmed for 1 h at 37°C to avoid crystals) in a non-CO2 incubator, with the plates sealed with Parafilm. The same protocol was applied to control and DMD cells. Bright-field images were acquired for qualitative analysis.

### In vitro myogenic differentiation assay

Myogenic cells were seeded onto Matrigel-coated dishes (Corning, cat. #356237) at confluence in growth medium to promote attachment. The following day, the medium was replaced with high-glucose, phenol red-free DMEM supplemented with 2% horse serum, L-glutamine, sodium pyruvate and 1% penicillin-streptomycin, and cells were kept in culture for 7 to 12 days. Differentiation was quantified as the myotube fusion index, using a slightly modified version of the ViaFuse workflow (Hinkle et al, 2021) in Fiji, calculated as the percentage of nuclei within MyHC-positive myotubes (containing ≥ 2 nuclei) over the total number of nuclei.

### Telomere fluorescence in situ hybridisation (FISH)

Telomere FISH was adapted from Tichy et al (2017). Cells (5,000–10,000) were seeded on Matrigel- or collagen-coated 8-well slides (LabTek, Merck) in growth medium and allowed to attach overnight. Cells were washed with PBS, fixed in 4% paraformaldehyde for 15 min, washed and permeabilised with 0.5% Triton X-100 in PBS for 15 min, and treated with pre-boiled RNase A (100 µg/ml) for 30 min at 37°C. A heating block was pre-warmed at 86°C, and slides were dried on it. The hybridisation solution contained 60% formamide, 33% distilled water, 2% 1 M Tris pH 7.5, 5% 10× blocking reagent for nucleic acid hybridisation (Roche; not diluted to 1×) and the Cy3-labelled C-rich telomere PNA probe (50 µM, 1:300; Eurogentec, PN-TC050-005), and was pre-heated at 86°C for 5 min. The mixture was applied to the slides and incubated for 10 min at 86°C in the dark, and slides were then cooled in the dark for 4 h to overnight. Slides were washed at 56°C in 2×, 1× and 0.5× SSC with 0.1% Tween 20, rinsed in PBS and mounted with VectaShield containing DAPI (Vector Laboratories, cat. H-1000-10).

### FISH image acquisition and quantification

Images were acquired with a DeltaVision Elite wide-field epifluorescence microscope (GE Healthcare) equipped with an Olympus 60×/1.42 Plan Apo N oil-immersion objective. For each field, 15–32 Z-stacks with a step of 0.2 µm were collected to cover the lower and upper boundaries of the DAPI signal, in the DAPI and Cy3 channels. Exposure times were set at the beginning of each session to place all samples within the mid-intensity range and avoid saturation, and the same acquisition parameters were applied to all samples within an experiment. Images were collected without binning, and stacks were deconvolved and projected with SoftWoRx (GE Healthcare). At least 50 nuclei per sample were analysed. Imaging sessions for each comparison were performed in the same time window to minimise variability in signal intensity.

Images were quantified with Telometer (DeMarzo Lab), an ImageJ macro, as described in Tichy and Mourkioti (2021), in a semi-automated fashion. Channels were normalised by background selection, each nucleus was outlined as a region of interest, and the macro measured the signal of every visible telomere within each region.

The value used for the analysis was “TelomereSumSumToCellSumPercent”, the percentage of telomere signal normalised to DAPI intensity. Results are shown as frequency distributions of nuclei.

### Karyotype analysis

Late-passage control cells and late-passage hTERT-treated DMD cells were analysed for chromosomal aberrations (at least 20 metaphases per sample) by the Cytogenetics Laboratory of the hospital (OSR), as a diagnostic cytogenetic report. Cells were arrested in metaphase with KaryoMAX colcemid (10 µg/ml; Gibco, 15212012) for a variable time depending on their division potential, and chromosomes were collected and stained with quinacrine for banding analysis.

### Statistical analysis

Data are expressed as mean ± SEM or SD, as indicated in the figure legends. Differences between groups were evaluated by one-way ANOVA followed by the post-hoc test given in each legend (Šídák’s, Dunnett’s or Tukey’s), or by unpaired t-test with Welch’s correction where only two groups were compared (Fig. S10), and p < 0.05 was considered significant. For Q-FISH, nuclei were the unit of analysis; as each comparison derives from a single experiment per donor, significance refers to differences between nuclei within that experiment. Post-transfection proliferation curves (one culture per condition and population) and the RNA-seq analysis were described without further statistical testing. Data were analysed with GraphPad Prism 10.3.

## Acknowledgements

Part of this work was carried out in ALEMBIC, an advanced microscopy laboratory established by IRCCS Ospedale San Raffaele and Università Vita-Salute San Raffaele. We also thank FRACTAL, the flow cytometry resource of IRCCS Ospedale San Raffaele, for support with flow cytometry analysis, and the Cytogenetics Laboratory of IRCCS Ospedale San Raffaele for the karyotype analyses.

## Author contributions

L.B.: Conceptualization, Methodology, Investigation, Formal analysis, Visualization, Project administration, Writing – original draft. F.M.: Conceptualization, Methodology, Resources, Supervision, Writing – review & editing. G.C.: Conceptualization, Funding acquisition, Supervision, Writing – review & editing.

