## Supplementary material for "Transient telomerase mRNA delivery extends the proliferative capacity of patient-derived Duchenne muscular dystrophy myogenic progenitors"

Barone, L.[1], Mourkioti F[2] and Cossu G[1,3].

### Contents

**Figure S1.** Representative example of the gating strategy used to discriminate the cells.

**Figure S2.** Representative bright-field images of myogenic progenitors at different stages of replicative ageing.

**Figure S3.** SA-β-gal (X-gal) staining of healthy and DMD myogenic progenitors at late passages.

**Figure S4.** Positive controls of the TRAP assay.

**Figure S5.** Integrity of the in vitro transcribed hTERT mRNA.

**Figure S6.** Permissiveness of cells to mRNA transfection.

**Figure S7.** hTERT expression one week after transfection.

**Figure S8.** Ki-67 and EdU co-staining in DMD #2 myogenic cells after hTERT mRNA transfection.

**Figure S9.** Telomere length distribution including mock-transfected controls.

**Figure S10.** Fold change of the telomere signal in pooled nuclei after hTERT mRNA transfection.

**Figure S11.** Hallmark gene sets.

**Figure S12.** Reactome pathway-level results of CD56+ cells.

**Figure S13.** Reactome pathway-level results of CD56− cells.

**Table S1.** Percentage of Ki-67-positive nuclei at early and late passages.

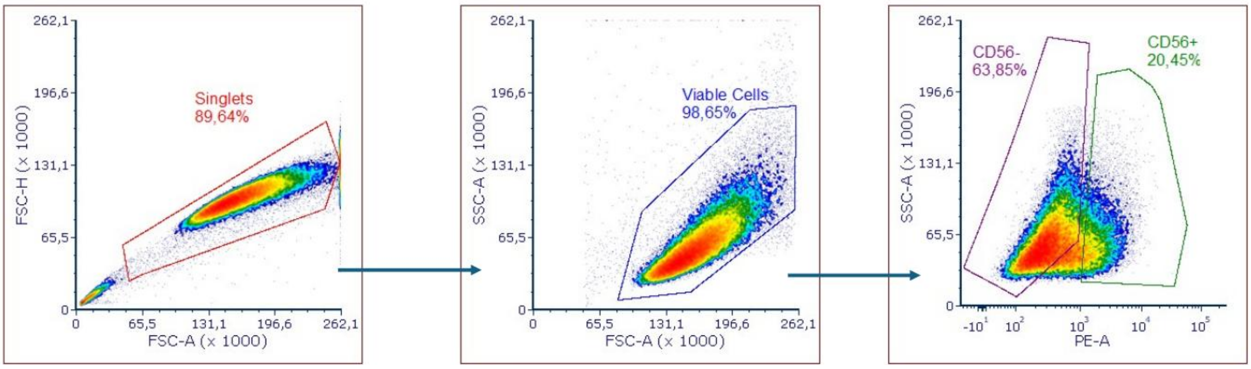


*Figure S1)* ***Representative example of the gating strategy used to discriminate the cells****. Cells were analysed for their physical parameters (FSC-H, FSC-A, SSC-A) to select a suitable fraction, and CD56+ and CD56− cells were then identified with a fluorochrome-conjugated antibody.*

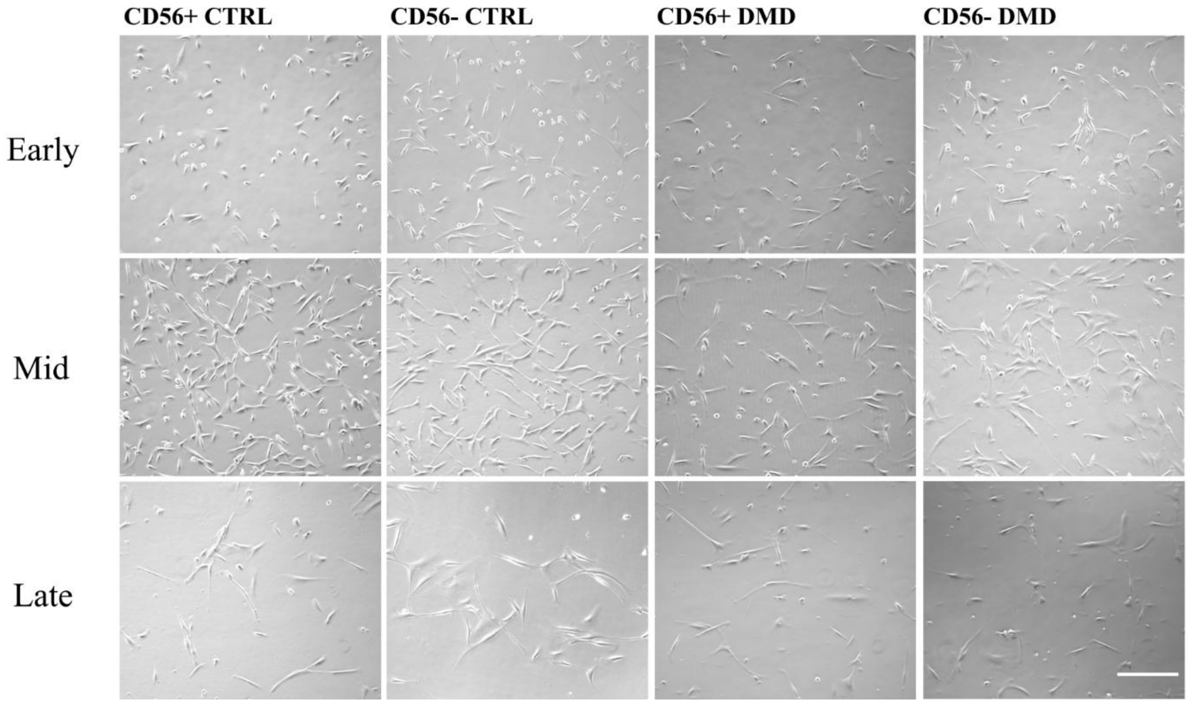


*Figure S2)* ***Representative bright-field images of myogenic progenitors at different stages of replicative ageing.*** *Representative images of CD56+ and CD56− populations from CTRL and DMD-derived cultures at different stages of culture. Scale bar: 500 µm.*

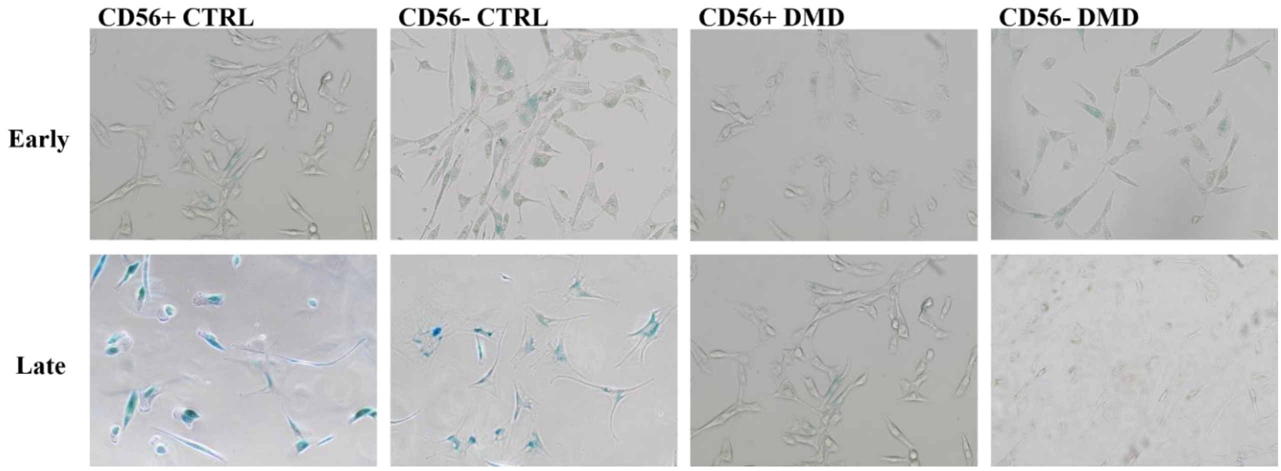


*Figure S3)* ***SA-β-gal (X-gal) staining of healthy and DMD myogenic progenitors at late passages.***
*Representative bright-field images of SA-β-gal-positive (blue) cells in CD56+ and CD56− populations from healthy and DMD donors. Staining was prominent in control cultures and weak or absent in DMD-derived cultures. SA-β-gal is not a specific marker of senescence.*

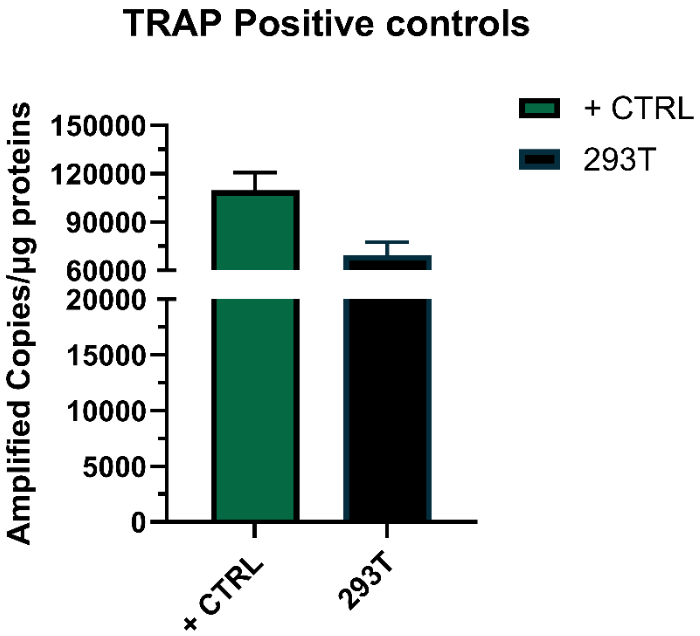


*Figure S4) Positive controls of the TRAP assay. A protein extract of HEK293T cells and the control cell pellet provided with the kit. The kit protocol includes only the sample named "+ CTRL", so HEK293T was used as an additional internal control.*

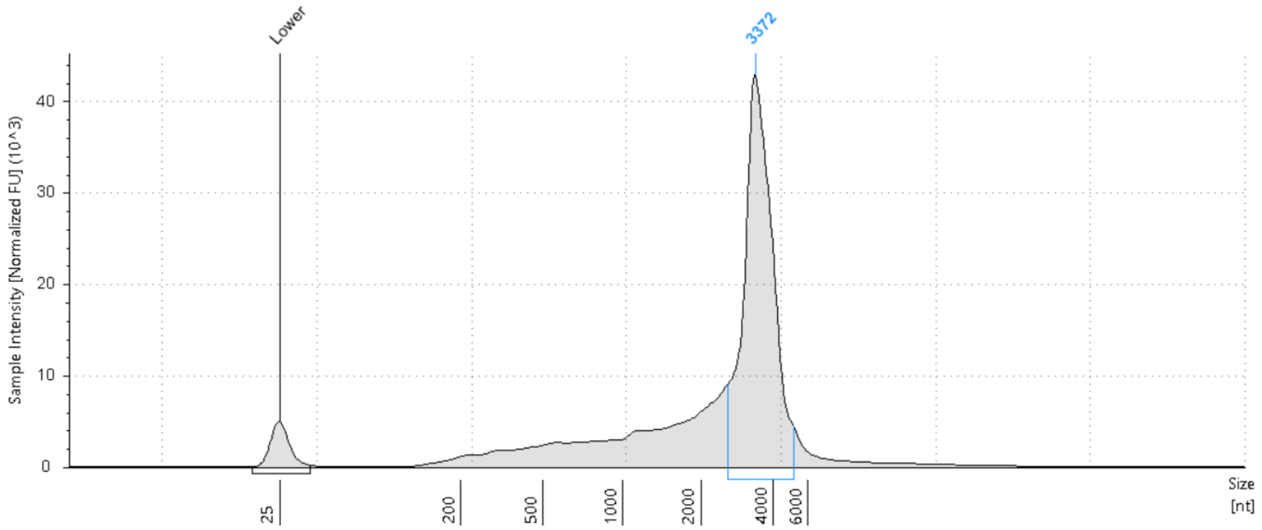


*Figure S5)* ***Integrity of the in vitro transcribed hTERT mRNA.*** *Capillary electrophoresis (TapeStation) of the purified transcript. The main peak migrates at ~3.4 kb (3,372 nt), consistent with the expected size of the transcript (~3.3 kb); the lower marker is at 25 nt.*

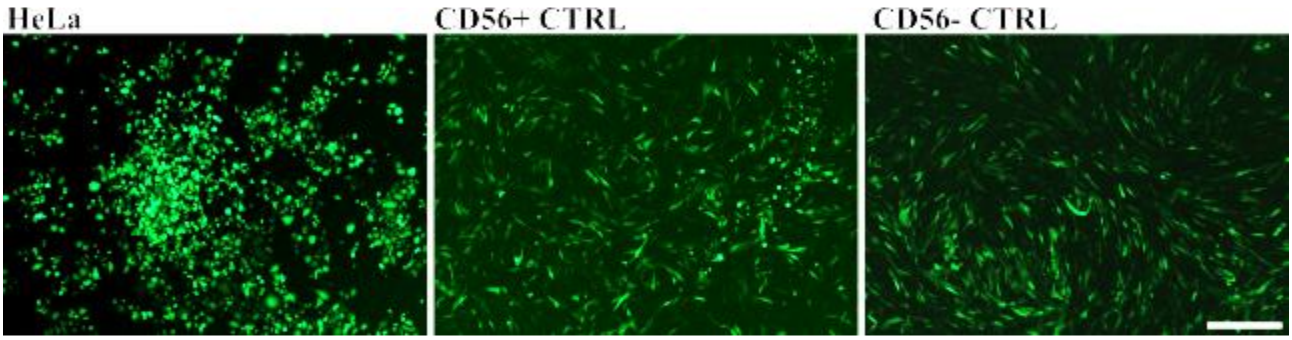


*Figure S6)* ***Permissiveness of cells to mRNA transfection.*** *HeLa cells and CD56+ and CD56− myogenic cells from the control donor were transfected with GFP-encoding mRNA (1 µg/ml) and imaged 24 h after transfection. GFP expression was detected in all three cell types. This was a qualitative assay and was not used to estimate transfection efficiency. Scale bar: 500 µm.*

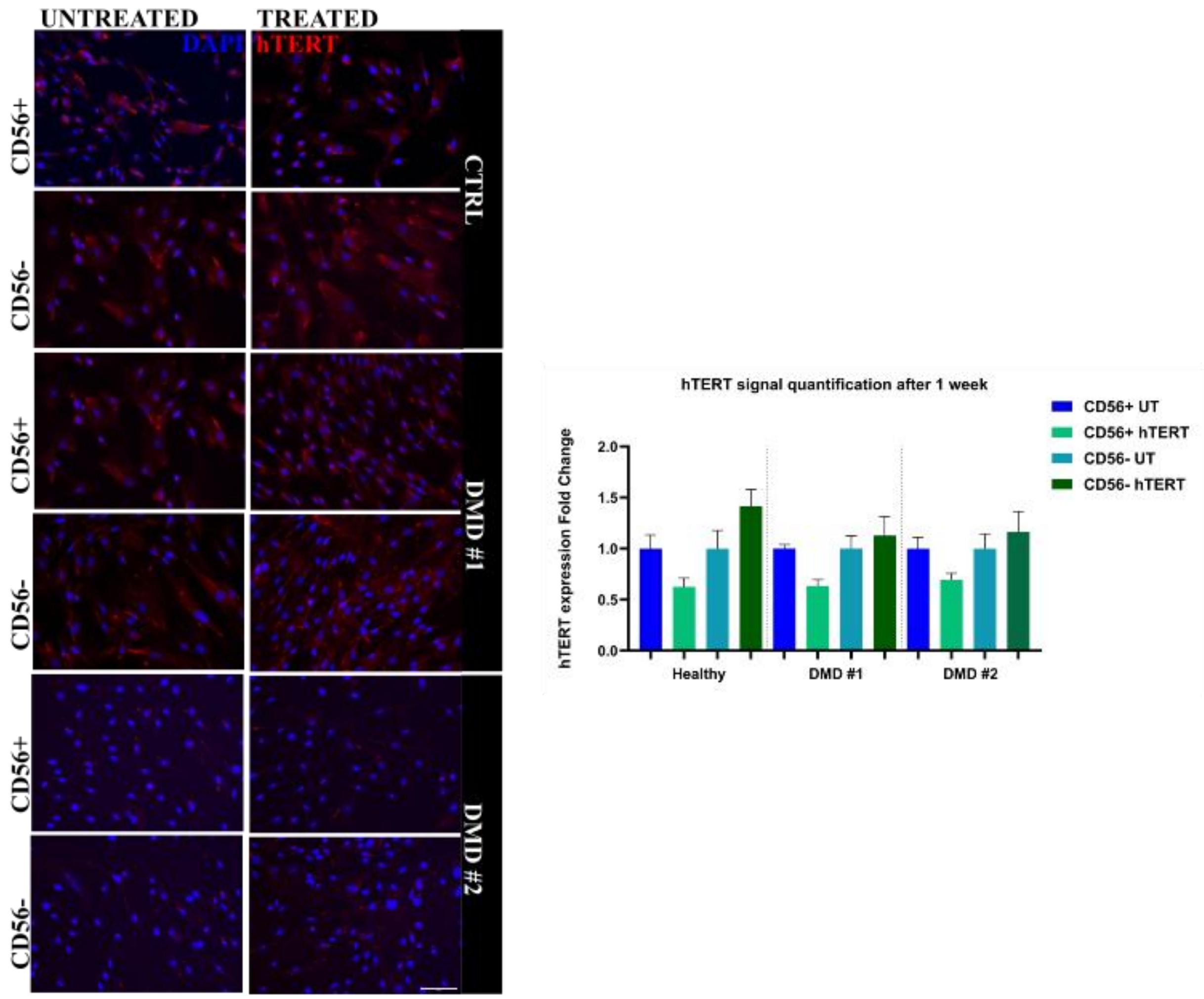


Figure S7) **hTERT expression one week after transfection.** Left: immunofluorescence for hTERT (red) and DAPI (blue) in untreated and hTERT mRNA-transfected CD56+ and CD56– cells from the control donor, DMD #1 and DMD #2, one week (two passages) after transfection. Right: hTERT signal (integrated density per nucleus, per field) expressed as fold change over the untreated sample of the same population (mean ± SEM of at least 5 independent fields). No statistically significant differences were observed between treated and untreated samples with one-way ANOVA, consistent with transient expression of the exogenous enzyme. Scale bar: 300 μm.

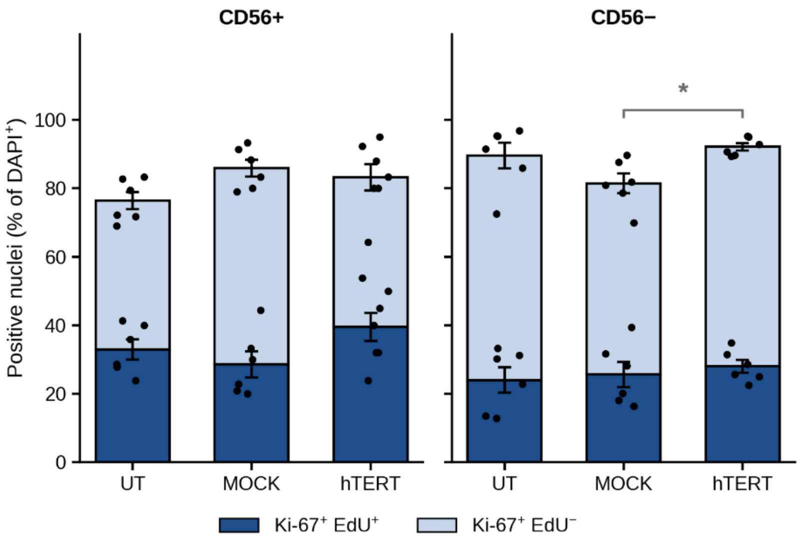


*Figure S8)* ***Ki-67 and EdU co-staining in DMD #2 myogenic cells after hTERT mRNA transfection.*** *Cells were analysed one week (two passages) after a 4 h transfection and pulsed with EdU for 2 h. Stacked bars show the percentage of nuclei positive for Ki-67 and EdU (dark) or Ki-67 only (light) in CD56+ and CD56− populations of untreated (UT), mock-transfected (MOCK) and hTERT mRNA-transfected cells. Percentages were calculated per field over total DAPI+ nuclei (CD56+: 222, 216 and 178 nuclei in UT, MOCK and hTERT; CD56−: 274, 514 and 417). Data are mean ± SEM of 6–7 fields per condition from a single experiment; dots represent individual fields. One-way ANOVA followed by Tukey's test was applied separately to Ki-67 and EdU; *p < 0.05 (CD56− Ki-67, MOCK vs hTERT). No other comparison was significant.*

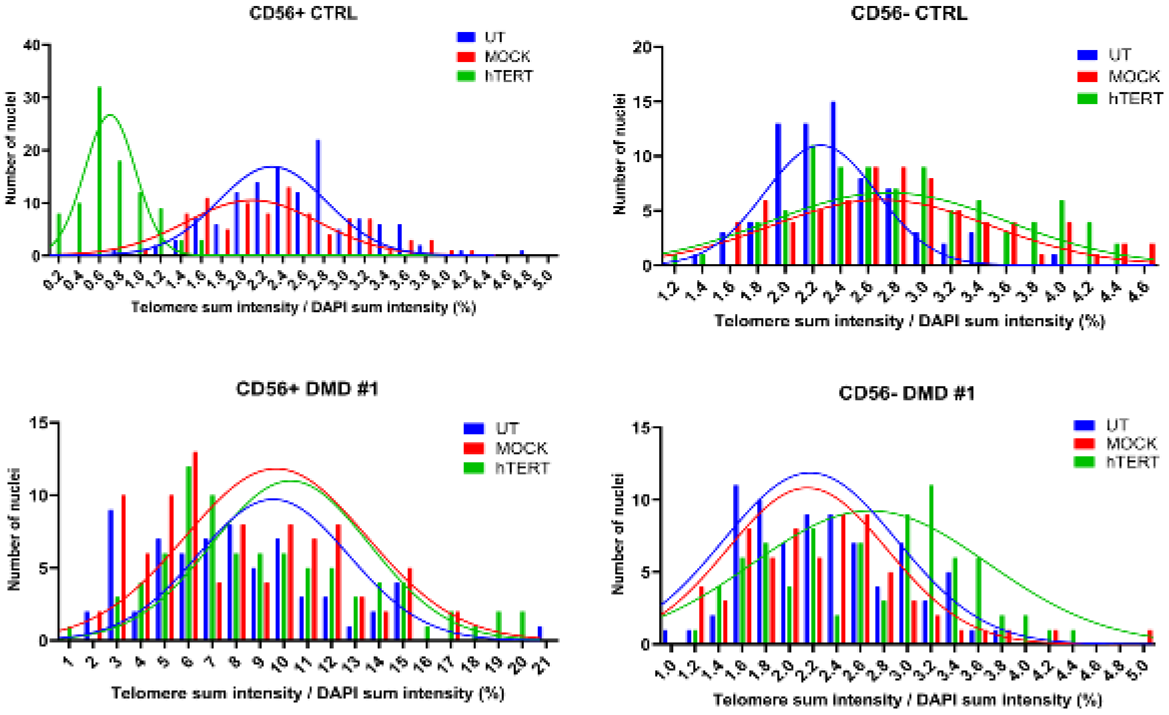


*Figure S9)* ***Telomere length distribution including mock-transfected controls.*** *Distribution of nuclei according to telomere sum intensity normalised to DAPI sum intensity (%) (Q-FISH) in untreated (UT, blue), mock-transfected (MOCK, red) and hTERT mRNA-transfected (hTERT, green) CD56+ and CD56− cells from the control donor and DMD #1, with Gaussian fits. These are the data shown in Fig. 4A with the addition of the MOCK condition, which was included only for these two donors. At least 50 nuclei were analysed per condition.*

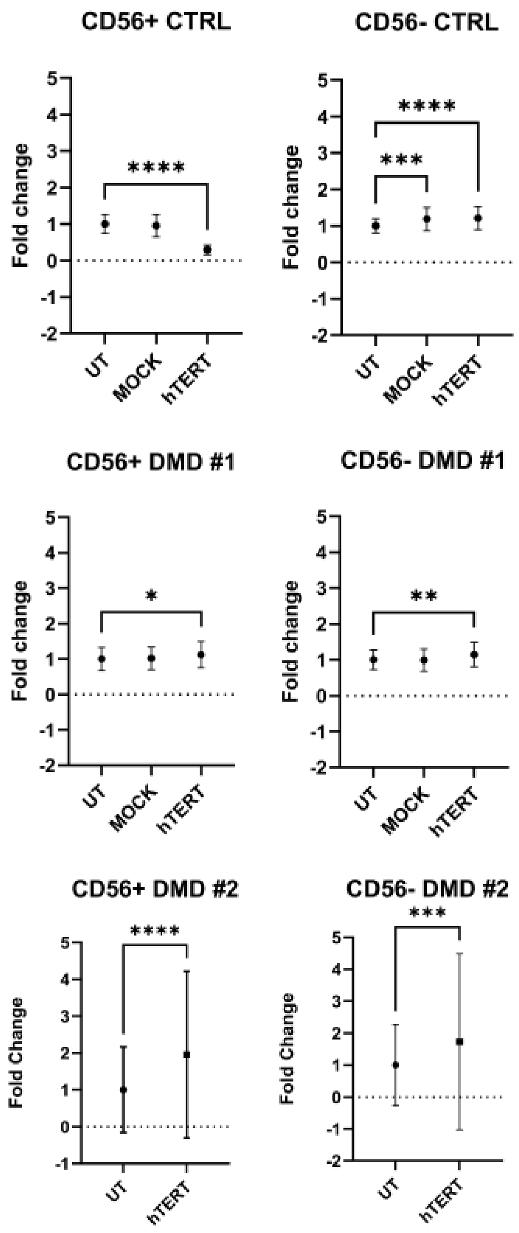


*Figure S10)* ***Fold change of the telomere signal in pooled nuclei after hTERT mRNA transfection.*** *Telomere sum intensity normalised to DAPI sum intensity (%) was measured per nucleus (Q-FISH) in CD56+ and CD56− cells from the control donor, DMD #1 and DMD #2. For each population, the values of all analysed nuclei were pooled and expressed as fold change over the mean of the untreated (UT) sample, to represent the shift of the treated distributions shown in Fig. 4A and Fig. S9. Mock-transfected controls (MOCK) were included for the control donor and DMD #1 only. Data are mean ± SD of at least 50 nuclei per condition, pooled for each population from a single experiment per donor. One-way ANOVA with Dunnett's multiple comparisons test versus UT (control donor and DMD #1); unpaired t-test with Welch's correction (DMD #2). *p < 0.05, **p < 0.01, ***p < 0.001, ****p < 0.0001.*

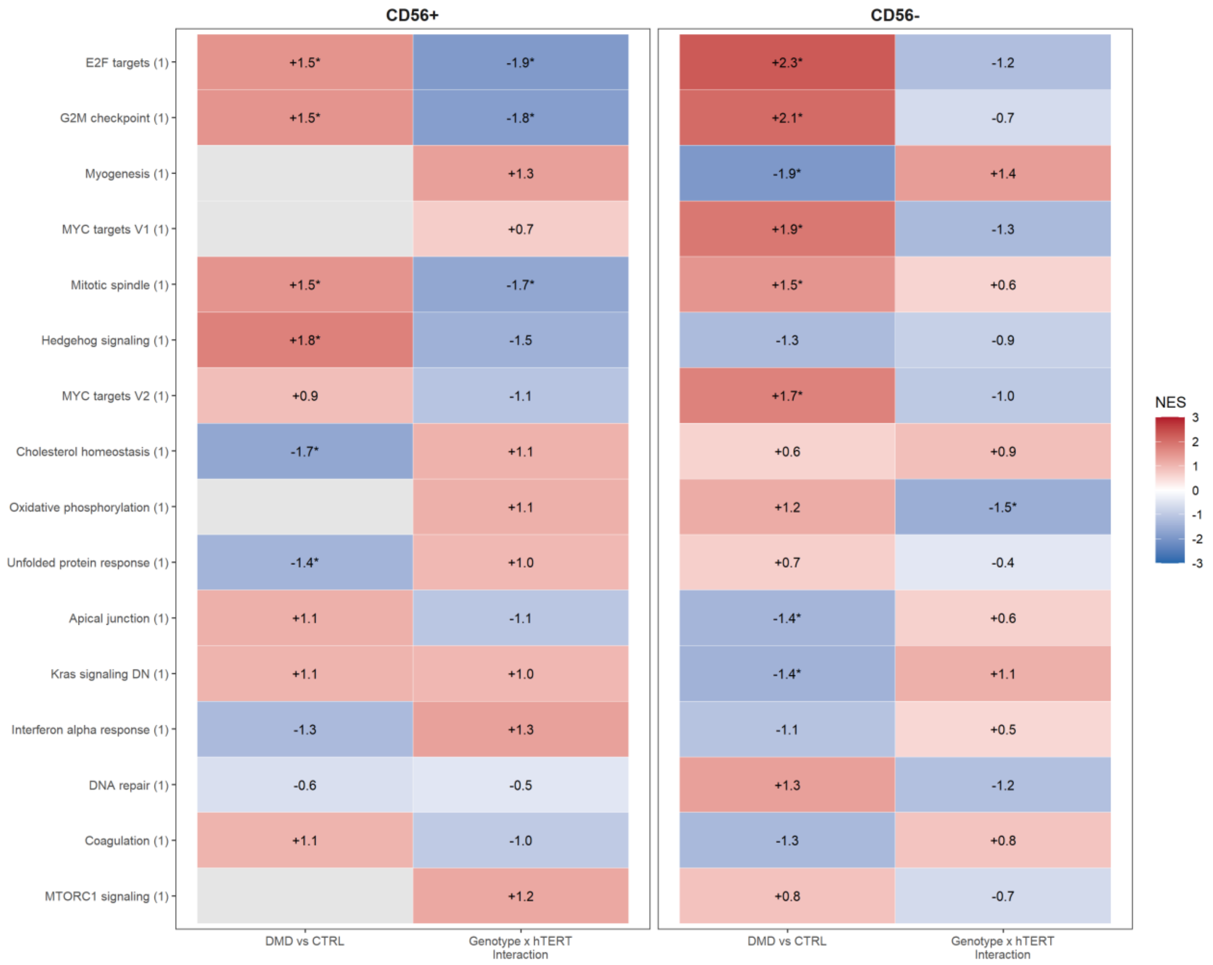


*Figure S11)* ***Hallmark gene sets.*** *Normalised enrichment score (NES) of Hallmark gene sets in CD56+ and CD56− cells for DMD vs control in untreated cells (DMD vs CTRL) and for the genotype × hTERT interaction. Each gene set is shown as a row (number of gene sets in parentheses). Red, positive NES; blue, negative NES; asterisks indicate padj < 0.05. One control and two DMD donors, no replicates; exploratory analysis.*

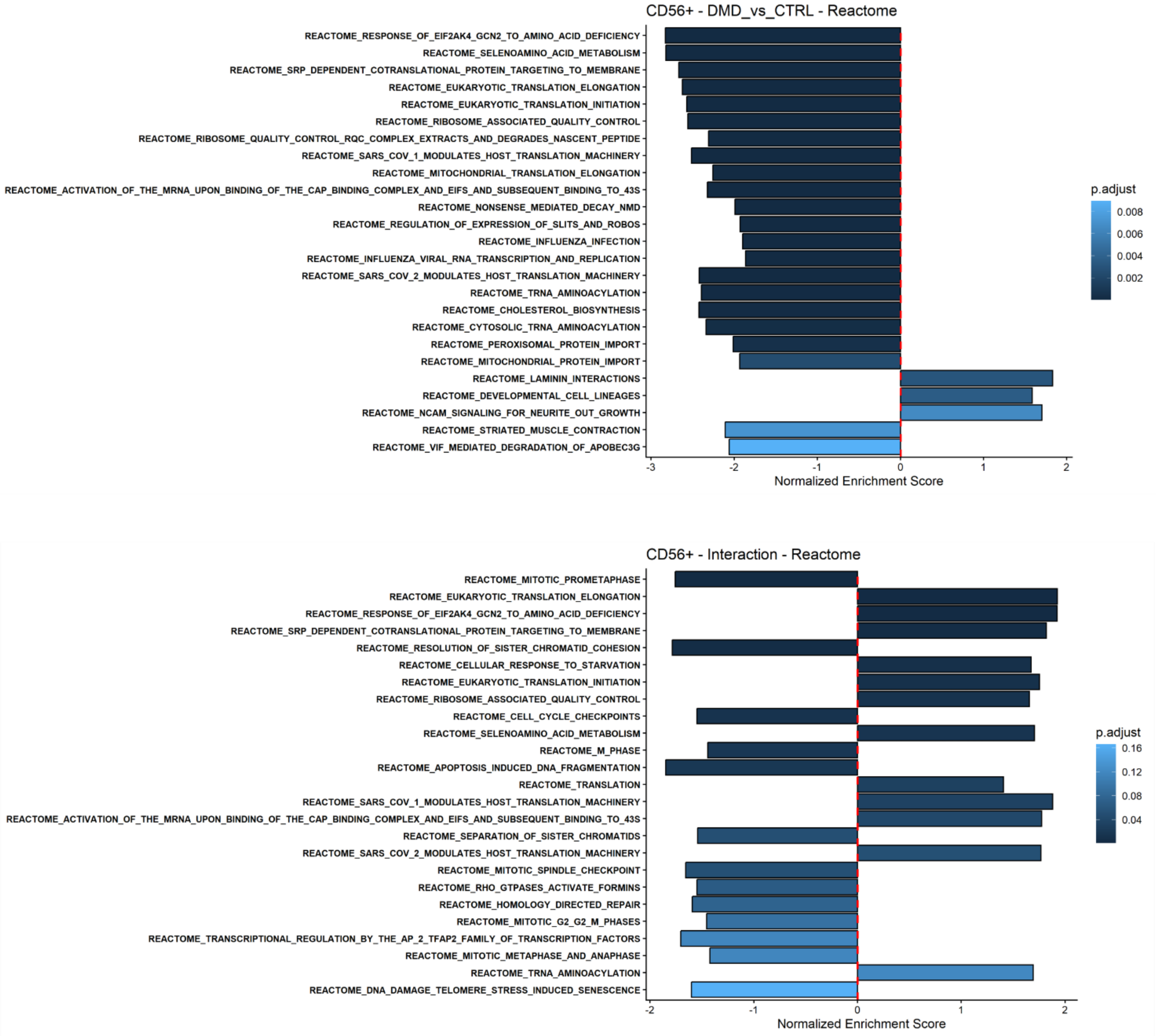


*Figure S12)* ***Reactome pathway-level results of CD56+ cells.*** *Normalised enrichment score (NES) of the 25 most significant Reactome pathways (padj < 0.25) for DMD vs control in untreated cells (DMD vs CTRL) and for the genotype × hTERT interaction, in CD56+ cells. Each pathway is shown as a bar, ordered by adjusted p value; the colour scale indicates the adjusted p value and the dashed line marks NES = 0. Pathway names are the original Reactome identifiers from MSigDB. One control and two DMD donors, no replicates; exploratory analysis.*

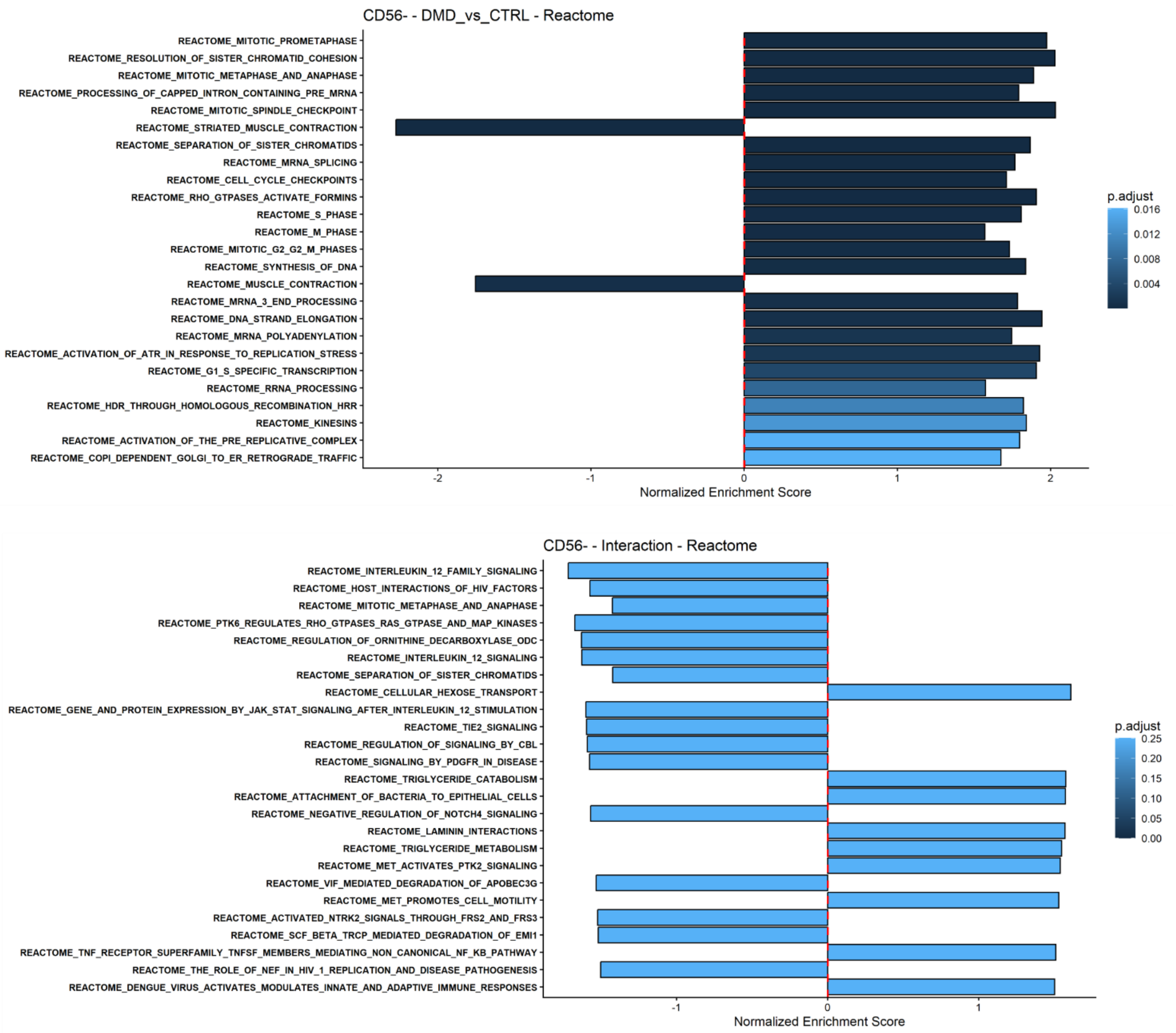


*Figure S13)* ***Reactome pathway-level results of CD56− cells.*** *Normalised enrichment score (NES) of the 25 most significant Reactome pathways (padj < 0.25) for DMD vs control in untreated cells (DMD vs CTRL) and for the genotype × hTERT interaction, in CD56− cells. Each pathway is shown as a bar, ordered by adjusted p value; the colour scale indicates the adjusted p value and the dashed line marks NES = 0. Pathway names are the original Reactome identifiers from MSigDB. One control and two DMD donors, no replicates; exploratory analysis.*

**Table S1.** Percentage of Ki-67-positive nuclei in CD56+ and CD56− cells from the control and the two DMD donors at early and late passages (data plotted in Fig. 1C).

| Population | Early passages (% Ki-67+ nuclei) | Late passages (% Ki-67+ nuclei) |
|---|---|---|
| Control CD56+ | 80.9 | 30.3 |
| Control CD56− | 55.1 | 38.6 |
| DMD #1 CD56+ | 55.4 | 10.7 |
| DMD #1 CD56− | 41.0 | 8.9 |
| DMD #2 CD56+ | 76.4 | 71.7 |
| DMD #2 CD56− | 89.6 | 78.5 |

*Values are means of at least three technical replicates per donor; at least 200 nuclei were analysed per condition. No statistical test was applied.*
